# Body size affects specialization and modularity in the global resin foraging meta-network of stingless bees

**DOI:** 10.1101/2023.04.10.536263

**Authors:** Daniel Yudi Miyahara Nakamura, Sheina Koffler, Tiago Mauricio Francoy

## Abstract

Bees are in global decline and specialized species might be the most vulnerable to extinctions. Bee conservation can be studied using interaction networks, whose relative importance of nodes might correlate with morphological traits. Specifically, body size could affect flying range and thus influence the set of plant-bee interactions. Although several studies have reported botanical sources of resins in stingless bees, resin foraging networks were rarely assembled. Here we aim to describe the global resin-foraging meta-network of stingless bees, identify the most specialized species, and test how body size influences modularity and specialization. We found a modular and nested structure, in which some modules exhibit significant differences in body size and specialization. *Melipona beecheii* is the most specialized stingless bee in collecting resins. Body size is positively correlated with specialization, in which larger bees are more specialized to collect resins from a subset of plants, possibly because larger bees with broader flying ranges avoid competition by collecting less disputed resources. Our results demonstrate how resin collection can be analyzed in a meta-network framework to test ecological hypotheses and identify specialized species as candidate priorities for the conservation of stingless bees.

## INTRODUCTION

Networks are widely employed to model ecological systems, in which nodes represent biological units and links depict interactions between nodes (Guimarães Jr. 2020). The number of possible topological configurations increases exponentially with the number of nodes but structured rather than random assemblies are repeatedly found in ecological systems (Olesen et al. 2007; Guimarães Jr. 2020). For example, two often emerged patterns are nestedness, wherein species with fewer interactions form a subset of interactions to species with more interactions (Bascompte et al. 2003), and modularity, in which species form more interactions among themselves within a module but fewer interactions with species from other modules (Jordano 1987; Dicks et al. 2002; Olesen et al. 2007; Fortuna et al. 2010). Moreover, interactions might exhibit phylogenetic signal, where closely related species share similar traits and tend to interact with a set of similar species due to a trophic niche conservatism (Wiens and Graham 2005; Rezende et al. 2007; Olalla-Tárraga et al. 2017; Corro et al. 2021).

Meta-networks comprise biotic interactions across different studies, that is, interactions between species that do not necessarily co-occur spatially or temporally in local populations (Heilmann-Clausen et al. 2017; Araujo et al. 2018; Emer et al. 2018) but may indirectly affect each other through a third species (González et al. 2018). Meta-networks may thus help understand mechanistic and evolutionary aspects of community organization, niche breadth, and the role of specialization in species interactions, as well as the impact of species loss on community stability (Blüthgen 2010). This approach could be useful for the conservation of bees since several studies have indicated declining trends in their diversity (Gallai et al. 2009; Garibaldi et al. 2009; Lever et al. 2014). However, except for a few studies (e.g. Leonhardt et al. 2011), resin collection is poorly studied in phylogenetic and meta-network frameworks, despite their important role in bee ecology (Requier and Leonhardt 2020; Shanahan and Spivak 2021).

Resins are lipid-soluble mixtures of volatile and non-volatile phenolic compounds and terpenoids (Langenheim 2003). Owing to their anti-inflammatory, antifungal, antibacterial, and antiviral properties, plants secrete resins to immobilize and defend themselves against predators and pathogens, as well as disinfect wound sites (Shanahan and Spivak 2021). For the same reasons, resins are used by bees to build nests, defense, and health (Requier and Leonhardt 2020). Considering a resin foraging network, morphological factors could influence centrality metrics at the node level (i.e., the relative importance of each node to the structure of the resin foraging network; Jordán et al. 2007). Specifically, body size could influence the diversity of botanical sources in some plant–bee interactions because larger species are expected to have a broader flying range than small species (Araújo et al. 2004). Thus, body size could be positively correlated with centrality metrics at the node level, in which specialist species tend to be smaller than generalists (Smith et al. 2019).

Stingless bees (Meliponini) comprise the most diverse group of corbiculate bees with more than 600 known eusocial species (Rasmussen et al. 2017; Melo 2021; Roubik 2022) and several uses of resins in their perennial nests (Roubik 1989). In stingless bees, resins are used during nest construction, incorporated in the nest entrance to defense, and mixed with wax or soil to produce cerumen or geopropolis, respectively (Wille 1983; Roubik 1989; 2006; Shanahan and Spivak 2021). Resins account for high proportions of foraging flights in Meliponini (Roubik 1989; Lorenzon and Matrangolo 2005) and despite their importance, little is known about their plant sources. However, three main methods are currently employed to identify botanical resin origins: (i) chemical analyses of resins and propolis, which are compared with chemical profiles of resins from local plants (e.g. Walker and Crane 1987; Bankova et al. 2000; Drescher et al. 2019); (ii) fieldwork, recording, or other visual observations (e.g. Wallace and Lee 2010; Gastauer et al. 2011; Reyes-González and Zamudio 2020); (iii) palynological analyses from pollen residues in propolis (Barth 1998; Barth et al. 1999; Barth and Lutz 2003; Barth 2006). Despite data availability in literature, a resin foraging meta-network was never assembled.

Here we provide insights into the global structure of resin foraging meta-network in stingless bees relying on a systematic review of the literature. First, we describe the network structure by testing whether nestedness and modularity are present in the topology and whether modularity is associated with node-level variables (body size and centrality metrics). Second, we identify resin foraging specialist species of stingless bees. Third, we analyze how body size affects specialization in plant–bee interactions. We expect that larger species tend to be generalists due to a broader flying range that increases access to more diverse plant sources. Furthermore, as a sensibility analysis, we also tested whether the network structure is affected by different methods of botanical source determination (chemical, fieldwork, and palynological) and whether sampling effort bias specialization in our data.

## MATERIAL AND METHODS

### Dataset

We conducted a systematic literature search in the Web of Science and Scopus databases. Following a similar approach to that described by Grames et al. (2019; see also Koffler et al. 2021), we employed the following keywords and some of their combinations: ‘botanical source*’, ‘geopropolis’, ‘Meliponini’, ‘plant source*’, ‘propolis’, ‘resin*’, and ‘stingless bee*’. The literature search was last performed on 5 February 2022 on titles, abstracts, and keywords. As potentially eligible articles, we also considered citations and references from publications included in the previous step. Duplicates were removed with the R package ‘litsearchr’ (Grames et al. 2019). We extracted additional terms using co-occurrence network analysis, which were used in a final search to enrich our dataset (see details in Supplementary Fig. S1). To fulfill objective criteria of inclusion, studies must have: (i) identified plant and stingless bee taxa at family and species level, respectively; (ii) specified the plant as a resin source rather than pollen source; (iii) indicated how they infer the interaction (chemical profile, fieldwork, or palynological analysis; Fig. 1A). Since several studies are only able to identify plants at the family level, we decided to relax the taxonomic resolution of plants from species to family because otherwise a large amount of information would be lost. The sampling effort was calculated as the number of papers reporting resin foraging interactions for each bee species. All sources of resin foraging data are in Supplementary Table S1.

**Figure 1:**
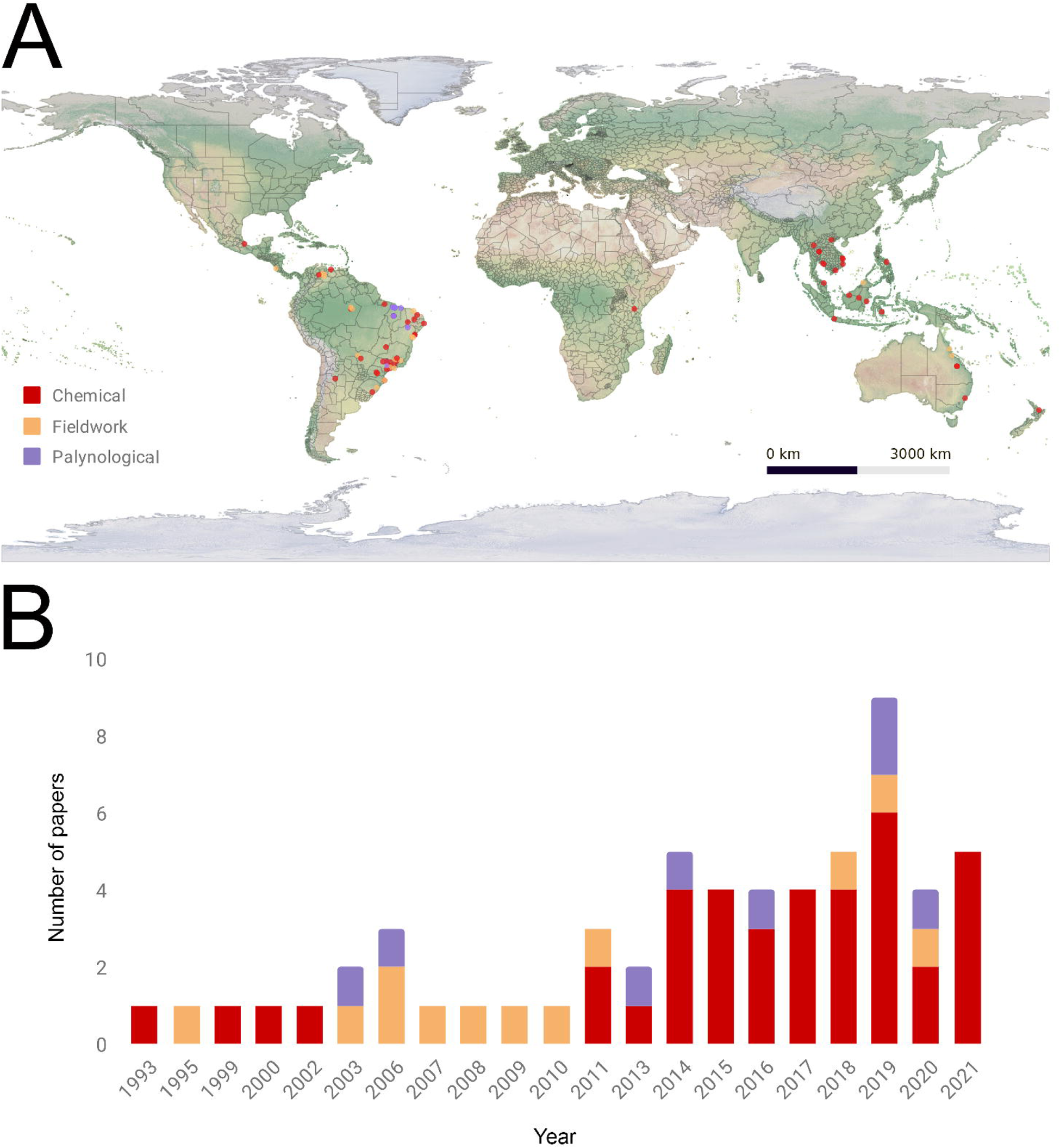
(A) Geographical distribution of studies reporting botanical origin of resins in stingless bees, separated by method employed for identification (chemical, fieldwork, or palynological). (B) Articles through years, separated by methods to determine botanical origins of resins.

While body size is a theoretical variable related to flight range, intertegular distance (ITD: the distance between the two insertion points of the wings; Cane 1987) was considered its proxy (Fig. 2). ITD was digitally measured with Zeiss ZEN at the Institute of Biosciences, University of São Paulo (IB-USP). Based on their availability, between three to ten specimens for each species were sampled from the Entomological Collection Paulo Nogueira Neto (CEPANN, IB-USP). ITD from species not available at CEPANN were searched in the literature (Supplementary Table S2).

**Figure 2:**
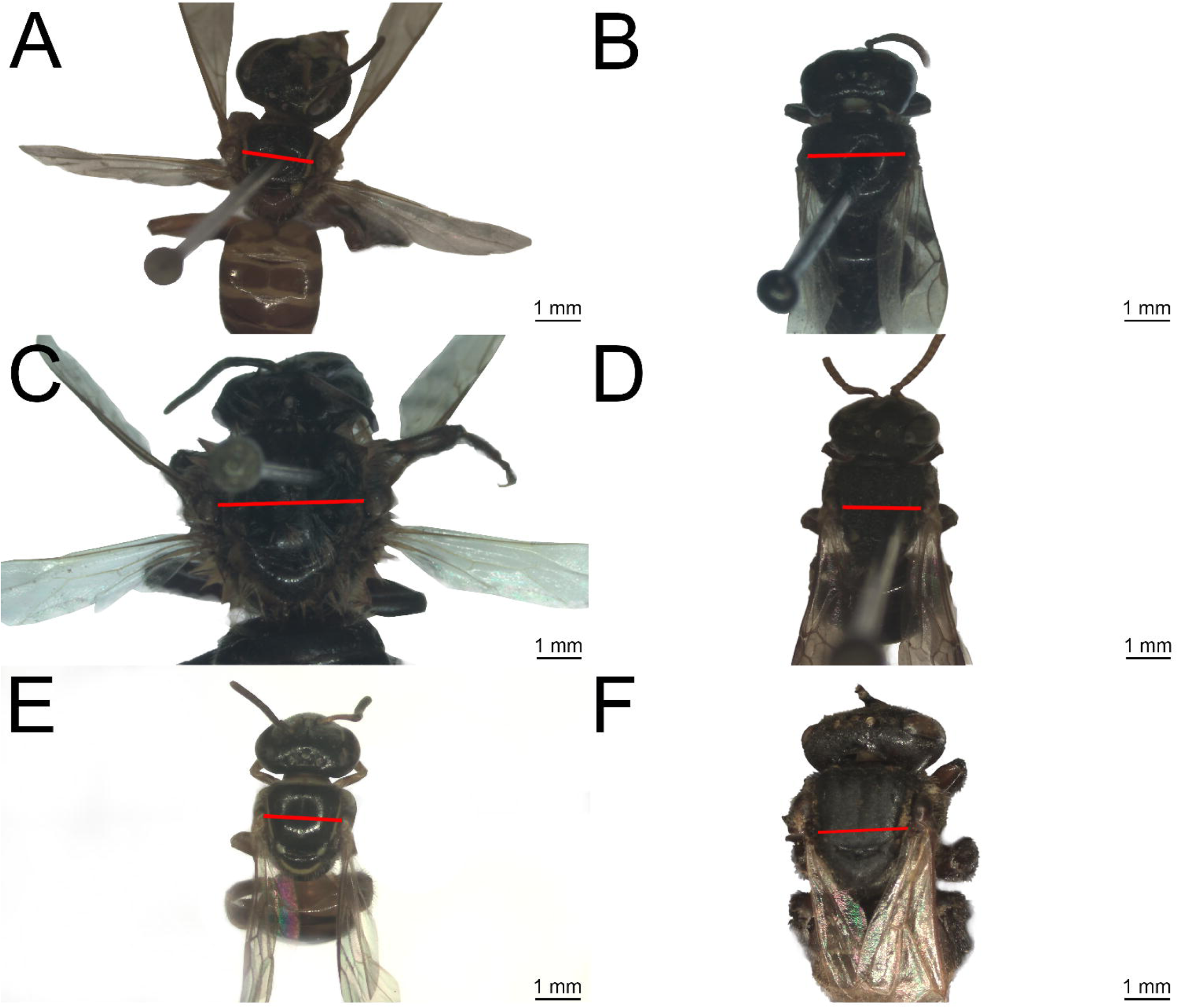
Examples of stingless bees measured in the present study, illustrating the diversity of intertegular distance (ITD used as proxy for body size). (A) *Frieseomelitta varia* (1.81 mm); (B) *Lestrimelitta limao* (2.33 mm); (C) *Melipona subnitida* (3.5 mm); (D) *Nannotrigona testaceicornis* (1.65 mm); (E) *Plebeia droryana* (1.51 mm); (F) *Scaptotrigona postica* (2.05 mm). Red lines indicate where ITD measurements were taken.

### Network analysis

The interactions dataset was organized as an adjacency matrix, with plant family represented in the rows and bee species in the columns, and the presence or absence of resin foraging computed in the cells. Hence, plants and bees are nodes, whereas the interactions between them are the edges of the bipartite network. Due to the variety of methods employed in the original studies to infer plant–bee interactions (chemical, fieldwork, and palynological analyses), we decided to use binary incidence matrices (presence vs absence) rather than weighted matrices, since biases could emerge if we mix interaction frequency data collected through distinct methods (Mello et al. 2019). In addition, binary data is indicated to assess fundamental niches rather than realized niches (Fründ et al. 2016; Jordano 2016). Our network was built at a global geographic scale, and hence its edges illustrate interactions between stingless bees and plant sources in the fundamental niche but not in their realized niches from specific local populations (Mello et al. 2019).

A set of centrality metrics are commonly used in the ecological literature to assess the relative importance of each node (plant and bee species) to the structure of a bipartite network (González et al. 2010). We employed four node-level metrics to characterize the relative importance of each species to the network structure. Bluthgen’s d specialization (hereafter, specialization d’) is measured as the selectiveness of the set of interactions performed by a bee concerning the interactions made by all other bee species in the network (Blüthgen et al. 2007). It means that a high specialization occurs when a bee species collects resins from a set of plant species that are not strongly collected by other bee species. Relative degree is the number of plant interactions with which a given bee species interacts scaled by the total number of plant species in the network (i.e., the potential number of interactions, which might be biologically interpreted as the fundamental niche breadth; Mello et al. 2015). Betweenness centrality is the proportion of shortest paths (i.e., geodesics) wherein a given bee species is present (Freeman 1977), which may be interpreted as the magnitude of a species in binding different guilds within the network (Mello et al. 2019). Closeness centrality is the average distance between a given bee species and all bee species in the network (Sabidussi 1966), which may be interpreted as a proxy for niche overlap (Mello et al. 2015). We reported the five most specialized bees in each network according to specialization d’, which can help to guide further studies and conservation programs (Raiol et al. 2021).

To describe the network structure, we estimated two network-level metrics using the ‘bipartite’ package. First, nestedness (NODF: range from 0 to 100; Almeida-Neto et al. 2008) indicates to what extent the links of low-degree nodes represent a subset of the links of high-degree nodes. Second, modularity (M) measures how much the network structure contains cohesive subgroups of nodes (modules) in which the density of interactions is higher within the same module than among modules. Both NODF and M were estimated for the global meta-network (NODF_global_, M_global_), as well as for meta-networks assembled from chemical (NODF_chemical_, M_chemical_), fieldwork (NODF_fieldwork_, M_fieldwork_), and palynological data (NODF_palynological_, M_palynological_). Statistical significance was estimated with null models using a Monte Carlo procedure (1000 simulated random matrices) for all analyses. Nestedness (NODF_null_) and modularity (M_null_) values were estimated for each null model, with Z-score calculated as Z = [observed value - mean (simulated values)] / σ(simulated values), following Mello et al. (2019). See the Appendix for details on the network- and node-level metrics.

### Statistical analyses

Because biological data from different species might be statistically non-independent due to a hierarchically structured phylogeny (Felsenstein 1985), we included the most comprehensive phylogeny of Meliponini in our statistical analyses (Quezada-Euán et al. 2019). The original tree includes 258 tips with representatives of all current genera. Before each analysis, we checked whether the terminal species from the tree matched the network-node species. Species with missing data were pruned from the tree. All taxonomic names followed the Integrated Taxonomic Information System (ITIS 2022).

To test the tendency of related species to exhibit similar node-level metrics and trait values under a model evolution, we estimated Pagel’s lambda (Pagel 1997, Pagel 1999). Statistical significance was calculated as the p-value of the likelihood ratio test for Pagel’s lambda using the *phylosig* function in ‘pythools’ (Revell 2012). We tested if ITD and centrality fitted three evolutionary models: (I) Brownian Motion (BM) assumes that traits change randomly and thus the correlation among trait values is proportional to the extent of shared ancestry for pairs of species (Felsenstein 1973); (II) Ornstein-Uhlenbeck (OU) is used to model trait evolution with the tendency towards an optimum value (e.g. under stabilizing selection; Butler and King 2004); and (III) Early-Burst (EB), in which most changes are concentrated towards the base of the tree (e.g. under adaptive radiation; Harmon et al. 2010). The Akaike Information Criterion corrected for small sample size (AICc) was performed with *fitContinuous* function in ‘geiger’ (Harmon et al. 2008). As a rule of thumb, if the difference between a model and the model with the minimum AICc is higher than two (ΔAIC > 2), the model with the lowest AICc is accepted as the most plausible (Burnham and Anderson 2002). To visualize ancestral states reconstructions, we employed the *contMap* function in ‘pythools’.

Phylogenetic Generalized Least Squares (PGLS) were performed using the function *gls* in ‘pythools’, with the error structure following the evolutionary model selected by ΔAIC. For each response variable (i.e., specialization d’, relative degree, betweenness centrality, and closeness centrality), we ran a PGLS model with log ITD as a fixed factor and contrasted it with a null model. These centrality metrics were analyzed separately as they were not correlated (Table S4). Additionally, we ran linear models including both log ITD and sampling effort as predictors. Sampling effort was included because less and more studied bee species could artificially produce lower and higher centrality metric values, respectively (Campião et al. 2015). In this case, linear models were employed rather than PGLS models because the sampling effort is not phylogenetically structured. To test differences of ITD and Bluthgen’s d specialization among modules, phylogenetic ANOVA was performed using the function *phylANOVA* in ‘phytools’, whereas Tukey’s test was run as a *post-hoc* analysis to detect which pair of modules exhibited significant differences.

## RESULTS

The naive search from Scopus and Web of Science resulted in 192 and 75 papers, respectively. After merging data (n = 267), 54 papers were deduplicated. The keyword co-occurrence network suggested the addition of the terms “botanical origin”, “plant origin”, and “plant resin”, which expanded the dataset from 213 to 243 papers. After screening papers reporting resin foraging interactions and plant and bee identifications, 183 papers were removed. Our final dataset included 60 studies (Supplementary Fig. S1; Table S1). Most studies were located in Neotropical and Indo-Malayan-Australasian regions, whereas botanical sources of resins in Africa were less reported (Fig. 1A). Moreover, our systematic literature search suggests that the number of papers reporting resin foraging interactions in stingless bees is increasing, with recent papers mostly determining the botanical source of resins comparing the chemical profile of resins between local plants and bees (Fig. 1B).

We assembled a meta-network with 99 plants, 67 stingless bees, and 493 unique plant–bee interactions (Fig. 3A). The network-level analysis revealed that nestedness is moderate (NODF_global_ = 34.58; Fig. 3B), a higher value than expected from null models (mean NODF_null-global_ = 24.24; Z = 6.59; p < 0.001; Fig. 3C). While chemical identification of botanical sources resulted in a balanced number of plant and bee nodes represented in the meta-network (Fig. 3D), fieldwork data resulted in meta-networks with more nodes representing bees (Fig. 3E) and palynological data resulted in meta-networks with a high number of plant nodes (Fig. 3F). Meta-networks assembled from data separated by method of identification of botanical source presented different levels of nestedness: NODF_chemical_ = 21.99 (NODF_null-chemical_ = 18.01; Z = 8.31; p < 0.001; Figs. 3D and S2A), NODF_fieldwork_ = 16.92 (NODF_null-fieldwork_ = 16.18; Z = 0.33; p = 0.673; Figs. 3E and S2B), and NODF_palynological_ = 53.59 (NODF_null-palynological_ = 45.33; Z = 3.83; p < 0.001; Figs. 3F and S2C).

**Figure 3:**
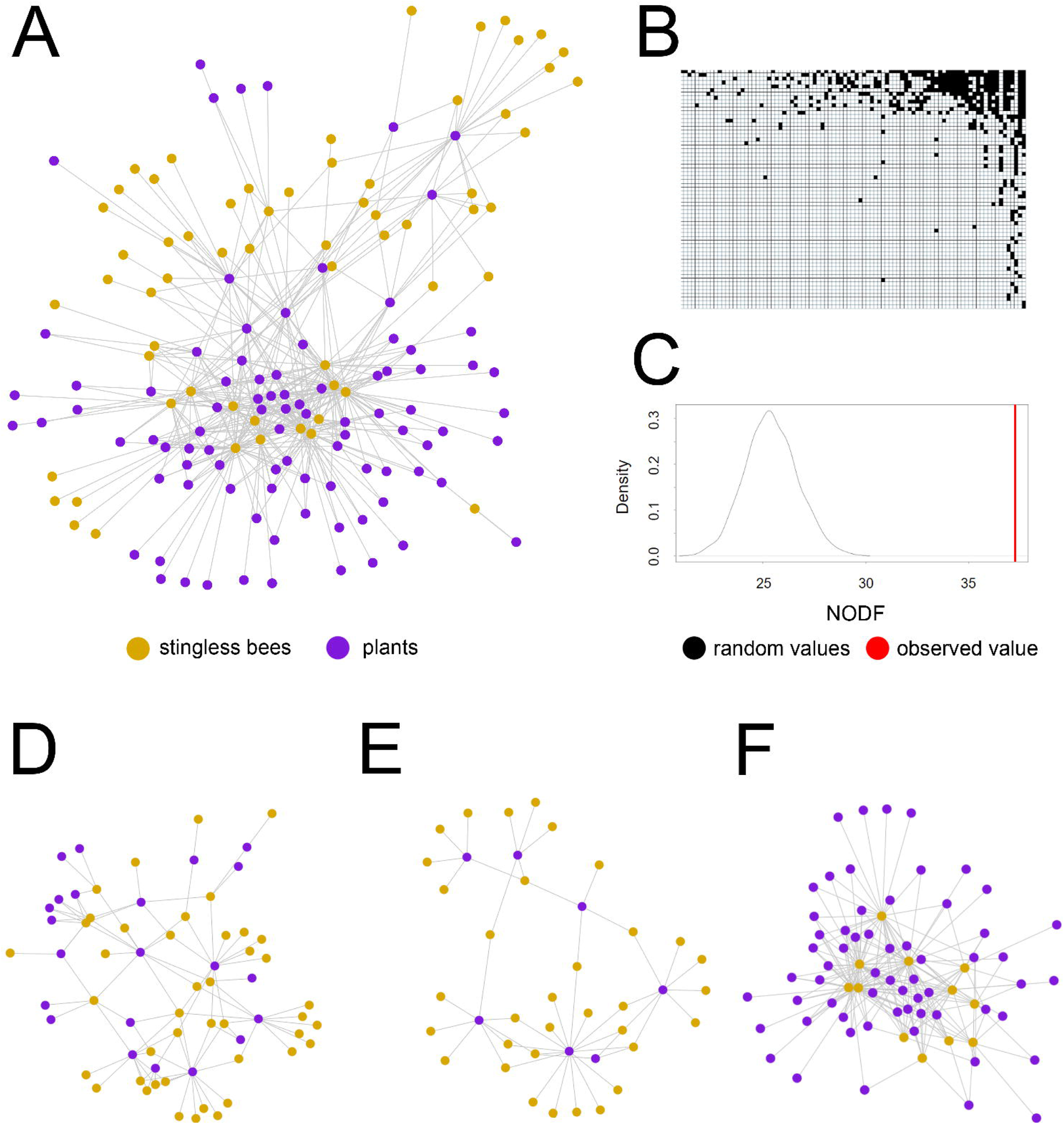
(A) Global resin foraging meta-network from interactions determined by all methods, (B) matrix showing the nested pattern in the global resin foraging meta-network of stingless bees, and (C) the distribution of random values generated by Monte Carlo null models against the observed NODF value. Resin foraging-meta-network separated by method of identification of botanical sources: (D) chemical profile, (E) fieldwork, and (F) palynological.

The modularity analysis of the global meta-network revealed five modules (M_global_ = 0.38; Fig. 4A and B; Supplementary Table S2), with a higher value than expected from Monte Carlo null models (M_null-global_ = 0.31; p < 0.001; Fig. 4C). Phylogenetic ANOVA revealed that body size (F = 7.215; p < 0.001; Fig. 4D) and centrality (F = 5.094; p = 0.002; Fig. 4E) were significantly different between modules, with Tukey’s test suggesting significant differences for body size between M1-M2 (p=0.006), M1-M4 (p = 0.003), M2-M3 (p = 0.026) and M3-M4 (p = 0.01; Supplementary Fig. S3A), whereas specialization d’ exhibited a significant difference between M1-M4 (p = 0.003; Supplementary Fig. S3B). Conversely, chemical, fieldwork, and palynological data presented modularity values of M_chemical_ = 0.56 (M_null-chemical_ = 0.50; Z = 3.48; p < 0.001; Supplementary Fig. S2D), M_fieldwork_ = 0.55 (M_null-fieldwork_ = 0.61; Z = −2.43; p = 0.98; Supplementary Fig. S2E), M_palynological_ = 0.55 (M_null-palynological_ = 0.24; Z = 32.69; p < 0.001; Supplementary Fig. S2F).

**Figure 4:**
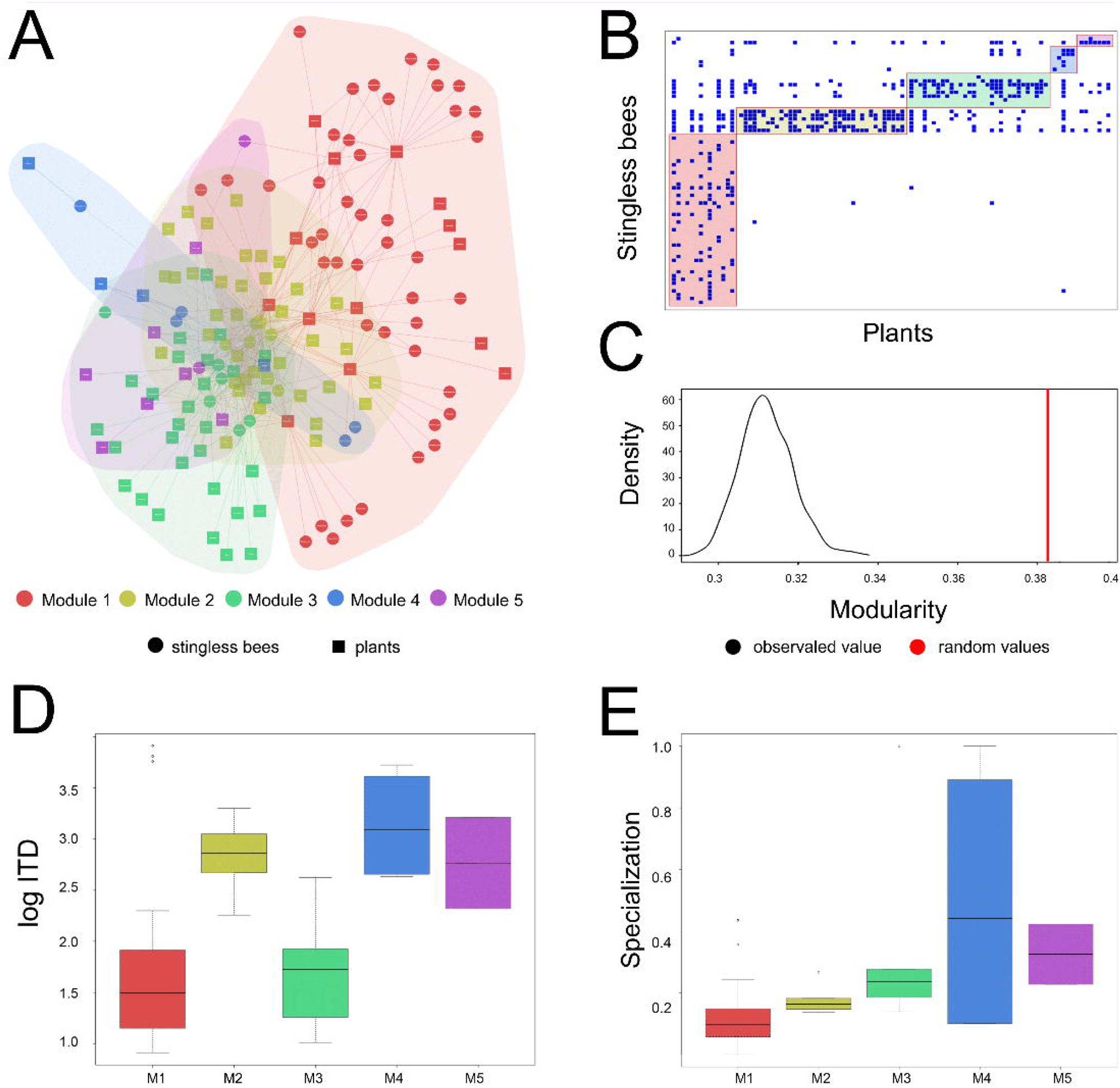
(A) Modularity in the global resin foraging meta-network of stingless bees, (B) matrix showing the modular pattern, and (C) the distribution of random values generated by Monte Carlo null models against the observed modularity value. Phylogenetic ANOVA between (D) modules and body size (ITD: intertegular distance; p = 0.00019), and (E) modules and specialization (p = 0.000707).

The node-level analysis suggested that *Melipona beecheii* and *Heterotrigona itama* are the most specialized and generalist species, respectively (see Supplementary Table S2 for specialization d’ values of all species). Phylogenetic signal was significant for body size (λ = 0.59; LR = 18.94; p < 0.001; Supplementary Fig. S4A) but not for specialization d’ (λ = 0.19; LR = 2.53; p = 0.11; Supplementary Fig. S4B), relative degree (λ = 0.03; LR = 0.16; p = 0.68), closeness centrality (λ = 6.61 × 10^−5^; LR = −0.01; p = 1.00), and betweenness centrality (λ = 0.01; LR = 0.01; p = 0.91). ΔAIC indicated that ITD and all centrality metrics are better fitted with the OU model (Supplementary Table S3). Model selection using AICc revealed body size as a predictor of specialization d’, but not for betweenness centrality, closeness centrality, and relative degree, in which null models were more plausible (Table 1). PGLS revealed a positive significant relationship between body size and specialization d’, indicating that larger bees are more specialized (Table 2; Fig. 5), whereas sampling effort did not affect specialization (Supplementary Table S4).

**Figure 5:**
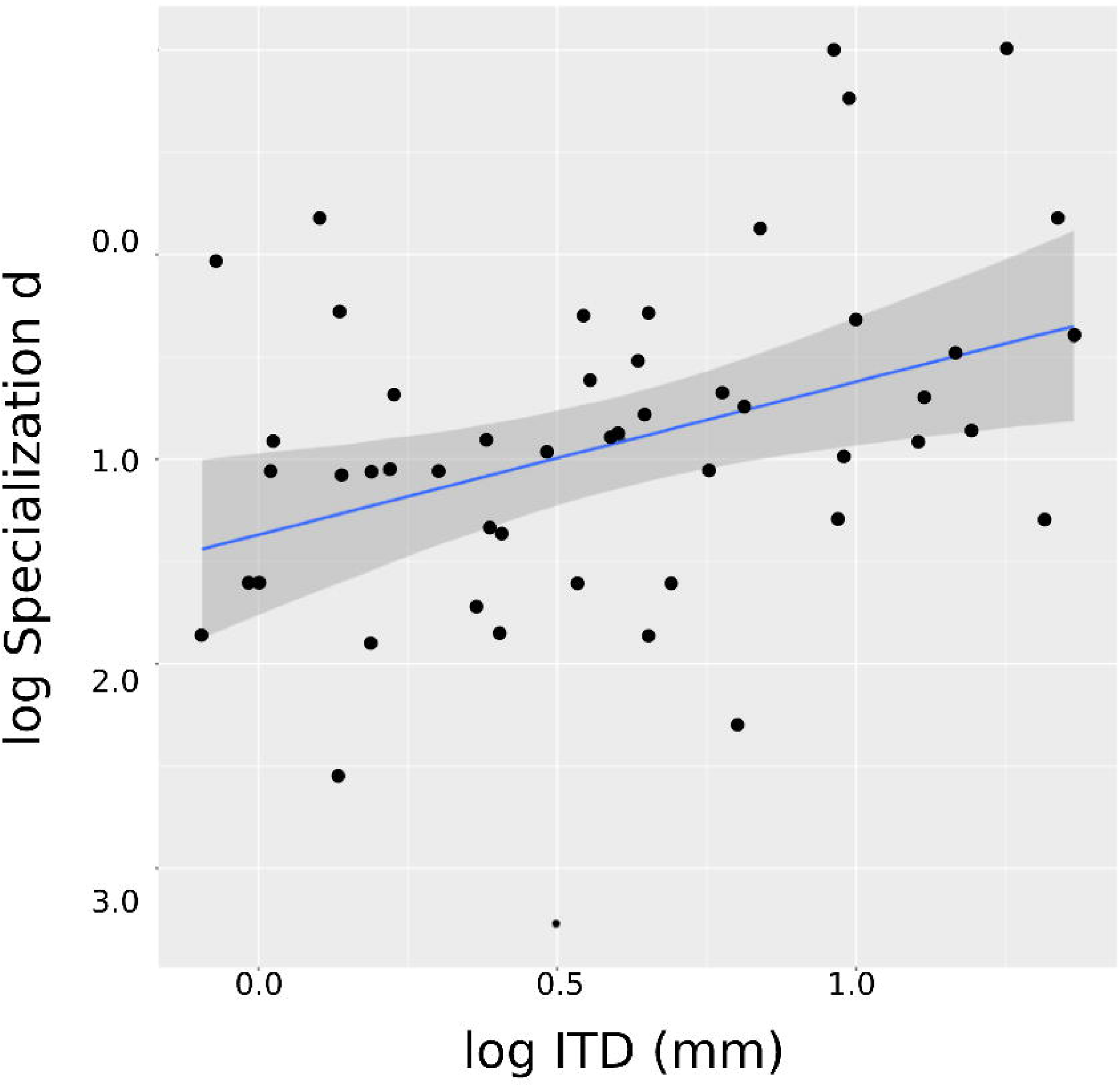
Scatter plot depicting the relationship between specialization d’ and ITD, according to PGLS following an Ornstein-Uhlenbeck error distribution.

**Table 1:**
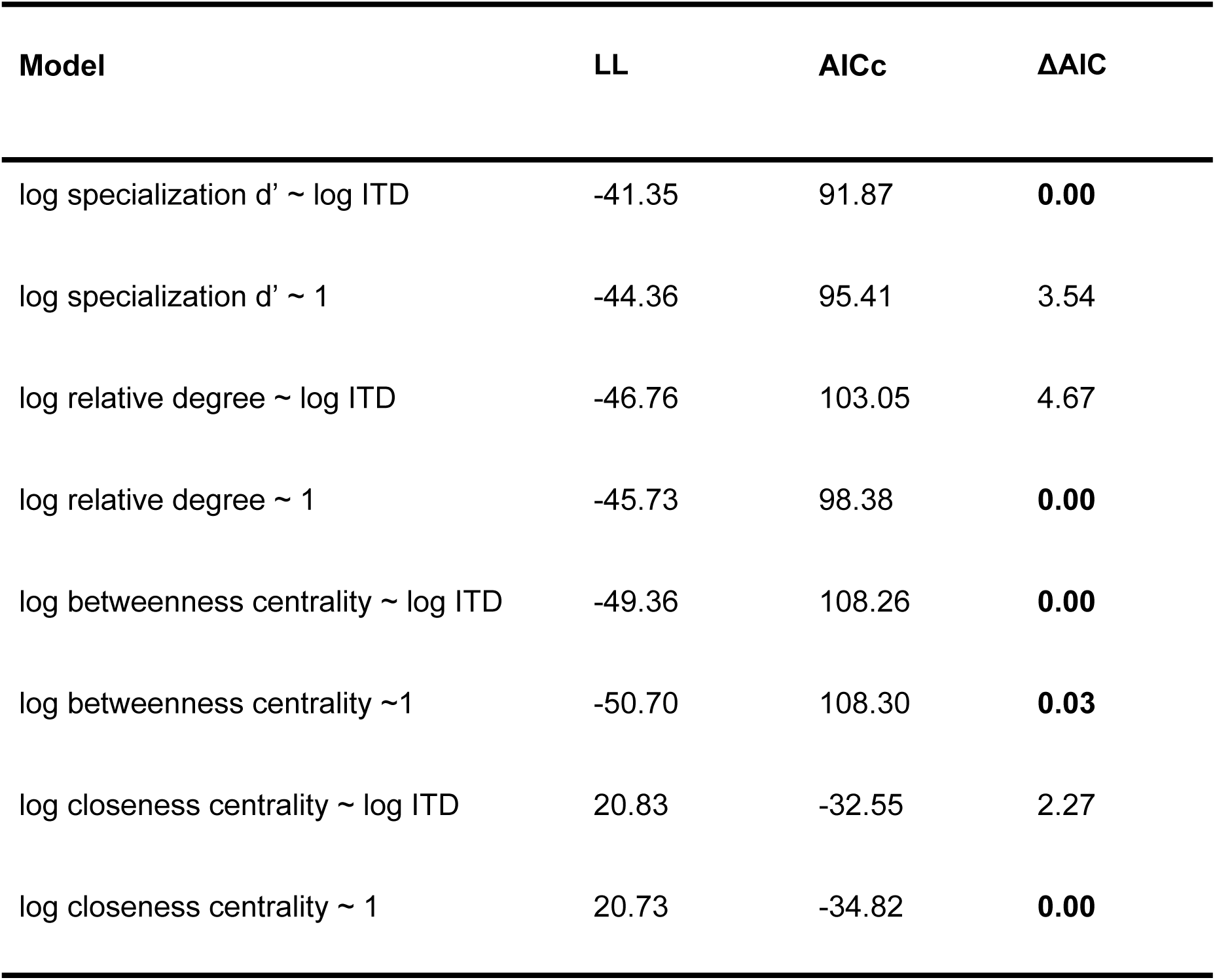
Model selection of PGLSs following an Ornstein-Uhlenbeck error structure. For each response variable, ΔAIC of best fitting models are in bold. The two PGLS models with betweenness centrality as response variable are equally plausible because their ΔAIC < 2. Abbreviations: LL = log-likelihood; AICc = AIC corrected for small sample size; ΔAIC = relative difference between the best model (= 0) and each other model.

**Table 2.** Best fitting PGLS model with Bluthgen’s specialization d’ as response variable and body size as predictor. Significant p-values are in bold.

| Coefficients | Estimate | Std. Error | t-value | p-value |
| --- | --- | --- | --- | --- |
| Intercept | -2.42 | 0.21 | -11.63 | **< 0.001** |
| log ITD | 0.71 | 0.29 | 2.48 | **< 0.001** |

## DISCUSSION

### Network-level metrics

The nestedness found in the global resin foraging meta-network of stingless bees indicates a system comprising less connected, peripheral bees with a high specificity to a few species of plants, whereas some subsets of more connected, central bees have a low specificity (Almeida-Neto et al. 2008; Cantor et al. 2017). Possible advantages of nestedness are the reduction in the mean number of interactions per species (Cantor et al. 2017), and hence a possible reduction in interspecific competition (Bastolla et al. 2009) and a potential increase in robustness against random extinctions (Memmott et al. 2004; Pocock et al. 2012). Although nestedness can simply emerge from the distribution of species abundances in neutral ecological networks (i.e., abundant species tend to interact with more nodes; Canard et al. 2012; 2014), life-history traits of plants (e.g., presence of spontaneous secretion of resins; Howard 1985) and bees (e.g., flight range; Inoue et al. 1985) also could affect resin collection when we consider niche-mediated processes to explain the nested structure.

The modularity found in this study indicates a non-random formation of semi-independent groups. Modularity might emerge either from niche overlap of similar species in the phylogeny (i.e., when closely related species with similar traits interact more with the same plants and thus are components from the same module; Lewinsohn et al. 2006; Olesen et al. 2007) or from convergence (i.e., when phylogenetically distant species are components from the same modules due to similar traits evolved through independent events; Thompson 2005). Our results indicate that modularity is partially shaped by body size, a trait associated with phylogeny. A possible explanation is that closely related stingless bees with similar size could present similar flight ranges (Casey et al. 1985; Byrne et al. 1988; Araújo et al. 2004). As such, species with similar flight ranges could exhibit niche overlap, interact with a high proportion of the same plant families, and thus be part of the same resin foraging module. For instance, small species with a short flight range might be restricted to collect resins from the same local plants (i.e., close to the nest), whereas large species with a broad flight range could access resins from distant plants. Furthermore, specialization d’ is significantly different between modules M1 and M4, indicating that species within M1 are more generalists, whilst species in module M4 exhibit a wide variation of specialization magnitudes.

### Node-level metrics

We tested whether body size affects centrality metrics in the resin foraging meta-network of stingless bees. However, contrary to our expectation, both PGLS (Table 2) and linear regressions suggest a positive relationship between body size and specialization d’, in which small bees tend to be generalists, whilst larger bees tend to be more specialists in resin collection. Moreover, our modularity analysis also seems to corroborate this pattern: the module with the lowest mean ITD (i.e., M1) is the same module with the lowest mean specialization; likewise, M4 exhibits the highest mean ITD and specialization d’.

In pollination interactions, the relationship between specialization and body size is controversial. Although our results contradict two previous studies in temperate environments (Leonhardt and Blüthgen 2012 in grasslands of Germany; Smith et al. 2019 in North American forests), it is in accordance with a tropical study (Raiol et al. 2021). In addition, a general pattern of positive relationships between size and degree was also found in distinct network types, suggesting a general mechanism (Chamberlain & Holland 2009). Resins are limiting resources for colonies of stingless bees (Howard 1985) and the idea that larger bees could monopolize resins and be dominant over smaller species (Johnson 1982; Slaa et al. 2003) favors the hypothesis that larger bees are generalists. However, Raiol et al. (2021) argue that warmer climates from tropical regions might support the evolution of specialists due to the high resource supply and the costs of its competition (Classen et al. 2020; Orr et al. 2021). In tropical environments, larger bees with broader flying ranges could specialize to forage resources untapped by smaller species with restricted flight ranges. Accordingly, large bees could even collect resins through bites that stimulate the secretion of exudates in plants (Schwarz 1948), whereas small bees may depend on the spontaneous secretion of resin or injury of the plant caused by other animals (Howard 1985; Leonhardt and Blüthgen 2009). As a result, because the cost of flying to distant resinous plants is possibly less than the cost of competition with small bees for local resinous plants, larger bees might avoid competition by specializing in the collection of less disputed resources and thus gaining fitness advantage (Raiol et al. 2021).

Two aspects of our tests must be considered. First, Ornstein-Uhlenbeck model is the best fitting model for all node-level metrics, which suggests that centrality (i.e., the relative importance of a node to the topology) evolved under stabilizing selection, but lineages possibly shifted optimum centrality values when new regimes were occupied (Butler & King 2004; O’Meara 2012). However, since the phylogenetic signal of node-level metrics is not significant, centrality of closely related species of bees is not necessarily similar. Second, because all node-level metrics partially measure centrality of nodes, some authors assume they are usually correlated and advocate in favor of using the first component of PCA as a centrality metric (e.g., Burin et al. 2021). Nevertheless, each metric provides different aspects of node importance (Appendix) and thus they are not necessarily correlated. Indeed, specialization d’ is the only variable not correlated with other node-level metrics (Supplementary Table S5), which reflects the results of specialization d’ being the only significant response variable of body size. Accordingly, specialization d’ is also the only node-level metric with no bias from the sampling effort (Table S3).

Generalist species are less vulnerable to losses of plants because their low specificity enables them to switch from one plant species to another, whereas the high specificity of specialists constrains this switch (Johnson and Steiner 2000; Biesmeijer et al. 2006; Steffan-Dewenter et al. 2006; Devictor et al. 2008; Klein 2011, but see Bommarco et al. 2010). For example, Jacquemin et al. (2020) used empirical plant-bee pollination data and demonstrated that specialist bees with few interactions tend to be lost over time. Thus, the calculation of specialization in resin foraging networks provides important information on priorities for conservation. Our results indicate that *Heterotrigona itama* is the most generalist species and *Melipona beecheii* – which collect resins from *Liquidambar styraciflua* (Altingiaceae) and *Populus nigra* (Salicaceae) in Mexico (Torres-González et al. 2016) – is the most specialized species. As such, the former is more central in the resin foraging meta-network, whereas the latter is more peripheral and thus could be more affected by potential extinctions of its botanical sources of resins. Thus, *L. styraciflua* and *P. nigra* should receive attention in conservation of *M. beecheii.* Future studies considering multiple interactions in the same network (i.e. multilayer networks; Mello *et al*. 2019) could be useful to better assess the relative importance of each species to bee conservation.

### Identification of botanical sources of resins

Resin collection might be hard to observe because this role is usually performed by a low relative number of worker bees (Simone-Finstrom and Spivak 2012) and it can occur high up in the trees (Crane 1990). Consequently, identifying botanical sources of these sticky products by direct observation (i.e., fieldwork) is challenging. In contrast, solutions have been proposed through palynological and chemical analyses over the past few decades. First, because propolis is resins mixed with salivary gland secretions and wax located on the nest floor or walls (Barth 1998; Bankova et al. 2000; Massaro et al. 2015; Shanahan and Spivak 2021), some authors argue that pollen grains stuck in propolis are suitable to characterize vegetation from which resins are collected (Barth 1998; Barth et al. 1999; Barth and Lutz 2003; Barth 2006). Second, plant sources of resins can be identified through mass spectrometry by comparing the chemical composition of resin loads carried by foragers with resins from local trees (e.g., Walker and Crane 1987; Bankova et al. 2000; Drescher et al. 2019). Due to the existence of different methods to identify botanical sources of resins, a reasonable question is whether these methods bias the calculation of network- and node-level metrics.

Regarding the calculation of network-level metrics, the assembly of resin foraging meta-networks revealed different patterns of nestedness but similar values of modularity across methods of identification of botanical sources. Specifically, NODF_palynological_ value is higher than NODF_chemical_ and NODF_fieldwork_. The chemical analysis produced balanced bipartite topologies (i.e., similar numbers of plants and bees), while meta-networks assembled from fieldwork and palynological data exhibited a relatively high number of bees and plants, respectively. This pattern emerges because fieldwork studies usually employ a focal observation of a single or a few species of plants while several species of bees visit them (e.g. Reyes-González and Zamudio [2020] observed a few scrubs of *Croton yucatanensis* while three species of stingless bees visited them), whereas palynological analysis is characterized by the identification of grains from several species of plants stuck in propolis or resin from a few species of bees (e.g. Barth [2006] found 44 families of plants in geopropolis samples from six species of Meliponini). Thus, the palynological meta-network exhibits a few species highly connected with plants, which explains its nestedness higher than chemical and fieldwork meta-networks.

Regarding the calculation of node-level metrics, our results revealed no evidence of an association between methods of identification of botanical sources and specialization d’ (Supplementary Fig. S5), which increases the evidence strength of our hypothesis tests using specialization d’ as a response variable. Here we chose a total evidence approach (i.e. using chemical, fieldwork, and palynological data) to assemble the global resin foraging meta-network, but future studies should experimentally test in local populations whether these methods bias the composition and abundance in resin foraging interaction data.

## Conclusion

Networks dealing with different interactions (e.g. pollination, nectavory, oil collection, and seed dispersal) have provided novel insights into macroecology, evolution, and conservation of bees (Memmott et al. 2004; Bezerra et al. 2009; Genini et al. 2010; Mello et al. 2013; Giannini et al. 2015; Kantsa et al. 2019; Mathiasson and Rehan 2020; Raiol et al. 2021). By incorporating interaction data of resin collection into a meta-network framework, we described its topological structure through network-level metrics, identified the most specialized species, and tested ecological hypotheses including body size as a predictor and node-level metrics as response variables. The resulting meta-network was modular and nested, with some modules significantly associated with body size and specialization. We revealed *Melipona beecheii* as the most specialized species, which could call attention to its conservation needs. Finally, PGLS revealed that body size was positively related to specialization, in which larger bee species are more specialized than smaller species.

## SUPPLEMENTARY INFORMATION

Appendix, supplementary tables, and figures are available in Supplementary Information.

## ACKNOWLEDGEMENTS

We thank Augusto Arruda for helping with measurements of intertegular distance, Isabel A. Santos for allowing us to access the Entomological Collection Paulo Nogueira Neto (CEPANN, IB-USP), and Marco A.R. Mello for his classes about ecological networks. We thank Marcelo Pompeo and Luis Carlos Souza of the Department of Ecology, Institute of Biosciences, University of São Paulo, for providing access to the ZEISS ZEN Microscopy.

## DATA AVAILABILITY STATEMENT

Body size data is provided in Supplementary Information. Binary incidence matrices of resin foraging interactions, phylogenetic tree, and R codes are available in the GitHub repository at https://github.com/danimelsz/resins

## AUTHOR CONTRIBUTIONS

Conceptualization: DYMN and SK. Methodology: DYMN and SK. Formal analysis and investigation: DYMN. Writing – original draft preparation: DYMN. Writing – review and editing: SK and TMF. Funding acquisition: TMF. Supervision: SK.

