## Supplementary for "Body size affects specialization and modularity in the global resin foraging meta-network of stingless bees"

Nakamura et al.

**Supplementary Tables** **2**

Table S1. 2

Table S2. 4

Table S3. 6

Table S4. 7

Table S5. 8

**Supplementary Figures** **9**

Figure S1. 9

Figure S2. 10

Figure S3. 11

Figure S4. 12

Figure S5. 13

**Appendix** **14**

**References** **17**

### Supplementary Tables

**Table S1.** Sources of resin foraging interactions, methods of determining botanical origins of resins, and number of bee species, plant species, plant genera, plant families, and interactions. Note that all bees were identified at species-level, whereas plant identification was heterogeneous across different studies.

| ID | Article | Method | Number of bee species | Number of plant species | Number of plant genera | Number of plant families | Number of interactions |
|---|---|---|---|---|---|---|---|
| 1 | Popova et al. (2021a) | Chemical | 4 | 1 | 1 | 1 | 4 |
| 2 | Thanh et al. (2021) | Chemical | 1 | 1 | 1 | 1 | 1 |
| 3 | Gabriel et al. (2021) | Chemical | 1 | 1 | 1 | 1 | 1 |
| 4 | Popova et al. (2021b) | Chemical | 1 | 1 | 1 | 1 | 1 |
| 5 | Oanh et al. (2021) | Chemical | 1 | 2 | 2 | 3 | 3 |
| 6 | Fikri et al. (2020) | Chemical | 2 | 1 | 1 | 1 | 2 |
| 7 | Maldonado et al. (2020) | Chemical | 2 | 12 | 8 | 7 | 17 |
| 8 | Reyes-González & Zamudio (2020) | Fieldwork | 3 | 1 | 1 | 1 | 3 |
| 9 | Silva et al. (2020) | Palynological | 1 | 0[1] | 20 | 15 | 25 |
| 10 | Sousa-Fontoura et al. (2020) | Chemical | 1 | 0[1] | 0[1] | 1 | 1 |
| 11 | Krahl et al. (2019) | Chemical | 1 | 1 | 1 | 1 | 1 |
| 12 | Ahmad et al. (2019) | Chemical | 1 | 1 | 1 | 1 | 1 |
| 13 | Lavinas et al. (2019) | Chemical | 1 | 1 | 1 | 1 | 1 |
| 14 | Pujirahayu et al. (2019) | Chemical | 1 | 1 | 1 | 1 | 1 |
| 15 | Jaapar et al. (2019) | Fieldwork | 11 | 4 | 3 | 2 | 34 |
| 16 | Georgieva et al. (2019) | Chemical | 1 | 4 | 4 | 4 | 4 |
| 17 | Annisava et al. (2019) | Chemical | 1 | 1 | 2 | 2 | 2 |
| 18 | Ribeiro et al. (2019) | Palynological | 4 | 14 | 108 | 46 | 277 |
| 19 | Negri & Silva (2019) | Palynological | 3 | 2 | 3 | 2 | 3 |
| 20 | Kraikongjit et al. (2018) | Chemical | 1 | 1 | 1 | 1 | 1 |
| 21 | Wang et al. (2018) | Fieldwork | 1 | 0[1] | 1 | 1 | 1 |
| 22 | Ishizu et al. (2018) | Chemical | 1 | 1 | 1 | 1 | 1 |
| 23 | Cisilotto et al. (2018) | Chemical | 2 | 1 | 1 | 1 | 2 |
| 24 | Souza et al. (2018) | Chemical | 1 | 0[1] | 1 | 1 | 1 |
| 25 | Pazin et al. (2017) | NA | 3 | 1 | 1 | 1 | 3 |
| 26 | Ferreira et al. (2017) | Chemical | 1 | 1 | 1 | 1 | 1 |
| 27 | Santos et al. (2017) | Chemical | 1 | 1 | 1 | 1 | 1 |
| 28 | Nguyen et al. (2017) | Chemical | 1 | 1 | 1 | 1 | 1 |
| 29 | Carneiro et al. (2016) | Chemical | 1 | 1 | 1 | 1 | 8[2] |
| 30 | Cunha et al. (2016) | Chemical | 1 | 0[1] | 2 | 2 | 2 |
| 31 | Torres-González et al. (2016) | Chemical | 1 | 2 | 2 | 2 | 2 |
| 32 | Ribeiro et al. (2016) | Palynological | 1 | 4 | 15 | 13 | 19 |
| 33 | Kustiawan et al. (2015) | Chemical | 1 | 2 | 2 | 1 | 2 |

| ID | Article | Method | Number of bee species | Number of plant species | Number of plant genera | Number of plant families | Number of interactions |
|---|---|---|---|---|---|---|---|
| 34 | Ragasa et al. (2015) | Chemical | 1 | 0[1] | 1 | 1 | 1 |
| 35 | Sanpa et al. (2015) | Chemical | 2 | 1 | 1 | 2 | 3 |
| 36 | Massaro et al. (2015) | Chemical | 1 | 16 | 9 | 6 | 16 |
| 37 | Massaro et al. (2014) | Chemical | 1 | 1 | 1 | 1 | 1 |
| 38 | Drescher et al. (2014) | Chemical | 1 | 3 | 4 | 3 | 4 |
| 39 | Souza et al. (2014) | Palynological | 1 | 3 | 75 | 34 | 83 |
| 40 | Dutra & de Barros (2014) | Chemical | 1 | 5 | 5 | 4 | 5 |
| 41 | Silva et al. (2014) | Chemical | 1 | 4 | 4 | 4 | 4 |
| 42 | Cunha et al. (2013) | Chemical | 1 | 0[1] | 0[1] | 1 | 1 |
| 43 | Barros et al. (2013) | Palynological | 1 | 4 | 31 | 25 | 38 |
| 44 | Leonhardt et al. (2011) | Chemical | 14 | 13 | 10 | 5 | 67 |
| 45 | Massaro et al. (2011) | Chemical | 1 | 1 | 1 | 1 | 1 |
| 46 | Gastauer et al. (2011) | Fieldwork | 4 | 2 | 3 | 3 | 6 |
| 47 | Wallace et al. (2010) | Fieldwork | 2 | 1 | 1 | 1 | 2 |
| 48 | Leonhardt et al. (2009) | Fieldwork | 11 | 8 | 7 | 4 | 27 |
| 49 | Freitas et al. (2008) | Fieldwork | 1 | 1 | 1 | 1 | 1 |
| 50 | Sawaya et al. (2007) | Fieldwork | 7 | 2 | 2 | 1 | 9 |
| 51 | Barth (2006) | Palynological | 6 | 7 | 31 | 44 | 147 |
| 52 | Sawaya et al. (2006) | Fieldwork | 1 | 1 | 1 | 1 | 1 |
| 53 | Bacelar-Lima et al. (2006) | Fieldwork | 2 | 1 | 1 | 1 | 2 |
| 54 | Moreno et al. (2003) | Fieldwork | 2 | 0[1] | 1 | 1 | 5[2] |
| 55 | Barth & Luz (2003) | Palynological | 3 | 0[1] | 23 | 44 | 107 |
| 56 | Patricio et al. (2002) | Chemical | 1 | 1 | 1 | 1 | 1 |
| 57 | Velikova et al. (2000) | Chemical | 4 | 1 | 1 | 1 | 4 |
| 58 | Lokvam & Braddock (1999) | Chemical | 1 | 1 | 1 | 1 | 1 |
| 59 | Wallace & Trueman (1995) | Fieldwork | 1 | 1 | 1 | 1 | 1 |
| 60 | Tomás-Barberán et al. (1993) | Chemical | 4 | 2 | 1 | 1 | 8 |

[1] Zero indicates that the authors were unable to identify the plant in that taxonomic level.

[2] The same plant-bee interaction was reported for different locations.

**Table S2.** Values of intertegular distance (ITD), biogeographical region, module, Bluthgen's specialization d', and references for ITD data. NA: values of specialization not applicable are present when a bee species is located in a small, isolated subnetwork rather than in the component with the highest proportion of nodes of the global meta-network.

| Bee species | ITD (mm) | Biogeographical region | Module | Specialization d' | Reference |
|---|---|---|---|---|---|
| *Axestotrigona ferruginea* | 1.925 | African | M3 | 0.275 | Quezada-Euán et al. (2019) |
| *Frieseomelitta silvestrii* | 1.010 | Neotropical | M3 | NA | Mayes et al. (2019) |
| *Frieseomelitta varia* | 1.740 | Neotropical | M3 | 0.199 | This study |
| *Geniotrigona thoracica* | 1.835 | Indo-Malayan-Australasian | M1 | 0.149 | Samsudin et al. (2018) |
| *Heterotrigona erythrogastra* | 1.150 | Indo-Malayan-Australasian | M1 | 0.125 | Samsudin et al. (2018) |
| *Heterotrigona itama* | 1.270 | Indo-Malayan-Australasian | M1 | 0.014 | Trianto and Purwanto (2020) |
| *Homotrigona apicalis* | 1.511 | Indo-Malayan-Australasian | M1 | 0.057 | Samsudin et al. (2018) |
| *Homotrigona binghami* | 1.495 | Indo-Malayan-Australasian | M1 | 0.029 | Samsudin et al. (2018) |
| *Homotrigona canifrons* | 1.805 | Indo-Malayan-Australasian | M1 | NA | Samsudin et al. (2018) |
| *Homotrigona fimbriata* | 1.925 | Indo-Malayan-Australasian | M1 | 0.057 | Samsudin et al. (2018) |
| *Homotrigona melanoleuca* | 1.588 | Indo-Malayan-Australasian | M1 | 0.065 | Samsudin et al. (2018) |
| *Lepidotrigona terminata* | 1.212 | Indo-Malayan-Australasian | NA | NA | Lee et al (2016) |
| *Lepidotrigona ventralis* | 1.440 | Indo-Malayan-Australasian | M1 | 0.029 | Sung et al. (2004) |
| *Lestrimelitta limao* | 2.181 | Neotropical | M3 | 0.186 | This study |
| *Lisotrigona cacciae* | 0.819 | Indo-Malayan-Australasian | M1 | 0.356 | Lee et al. (2016) |
| *Lisotrigona furva* | 1.027 | Indo-Malayan-Australasian | M1 | 0.148 | Lee et al. (2016) |
| *Melipona beecheii* | 2.681 | Neotropical | M4 | 0.782 | This study |
| *Melipona compressipes* | 3.725 | Neotropical | M4 | 0.101 | This study |
| *Melipona fasciculata* | 3.300 | Neotropical | M2 | 0.155 | Borges et al. (2020) |
| *Melipona favosa* | 2.633 | Neotropical | M4 | 0.101 | Quezada-Euán et al. (2019) |
| *Melipona flavolineata* | 2.671 | Neotropical | M2 | 0.137 | This study |
| *Melipona fuliginosa* | 3.810 | Neotropical | M1 | 0.436 | Mayes et al. (2019) |
| *Melipona mandacaia* | 2.708 | Neotropical | M2 | 0.266 | This study |
| *Melipona marginata* | 2.299 | Neotropical | M1 | NA | Quezada-Euán et al. (2019) |
| *Melipona orbignyi* | 3.500 | Neotropical | M4 | NA | Schwarz (1932) |
| *Melipona quadrifasciata* | 3.211 | Neotropical | M5 | 0.228 | This study |
| *Melipona scutellaris* | 3.912 | Neotropical | M1 | 0.244 | This study |
| *Melipona seminigra merrillae* | 3.020 | Neotropical | M2 | 0.147 | This study |
| *Melipona subnitida* | 3.050 | Neotropical | M2 | 0.182 | This study |
| *Nannotrigona testaceicornis* | 1.624 | Neotropical | M3 | 0.139 | This study |
| *Paratrigona anduzei* | 2.000 | Neotropical | M1 | 0.073 | Schwarz (1948) |
| *Plebeia droryana* | 1.476 | Neotropical | M1 | 0.096 | This study |
| *Plebeia emerina* | 1.241 | Neotropical | M1 | 0.127 | This study |
| *Plebeia frontalis* | 1.020 | Neotropical | M1 | 0.127 | Quezada-Euán et al. (2019) |
| *Plebeia lucii* | 1.000 | Neotropical | M1 | 0.073 | Teixeira and Campos (2005) |
| *Plebeia remota* | 1.865 | Neotropical | M1 | NA | This study |
| *Scaptotrigona bipunctata* | 2.320 | Neotropical | M5 | 0.424 | This study |

| Bee species | ITD | Biogeographical region | Module | Specialization d' | Reference |
|---|---|---|---|---|---|
| *Scaptotrigona depilis* | 1.710 | Neotropical | M1 | 0.073 | Mayes et al. (2019) |
| *Scaptotrigona postica* | 2.257 | Neotropical | M2 | 0.173 | This study |
| *Tetragona clavipes* | 1.886 | Neotropical | M1 | 0.219 | This study |
| *Tetragonisca angustula* | 1.257 | Neotropical | M3 | 0.185 | This study |
| *Tetragonisca fiebrigi* | 1.145 | Neotropical | M4 | 0.277 | This study |
| *Tetragonula biroi* | 1.210 | Indo-Malayan-Australasian | M1 | 0.127 | Trianto and Purwanto (2020) |
| *Tetragonula carbonaria* | 1.109 | Indo-Malayan-Australasian | M1 | 0.435 | Wittwer and Elgar (2018) |
| *Tetragonula collina* | 1.380 | Indo-Malayan-Australasian | M1 | 0.094 | Lee et al. (2016) |
| *Tetragonula fuscobalteata* | 1.006 | Indo-Malayan-Australasian | M1 | 0.057 | Lee et al. (2016) |
| *Tetragonula geissleri* | 1.915 | Indo-Malayan-Australasian | M1 | 0.169 | Samsudin et al. (2018) |
| *Tetragonula laeviceps* | 1.207 | Indo-Malayan-Australasian | M1 | 0.055 | This study |
| *Tetragonula pagdeni* | 0.981 | Indo-Malayan-Australasian | M1 | 0.073 | Lee et al. (2016) |
| *Tetragonula sapiens* | 3.760 | Indo-Malayan-Australasian | M1 | NA | Trianto and Purwanto (2020) |
| *Trigona corvina* | 2.125 | Neotropical | M1 | 0.127 | Schwarz (1948) |
| *Trigona fulviventris* | 1.360 | Neotropical | M1 | 0.128 | Quezada-Euán et al. (2019) |
| *Trigona recursa* | 1.724 | Neotropical | M3 | 0.272 | This study |
| *Trigona spinipes* | 2.224 | Neotropical | M1 | 0.037 | This study |
| *Trigona williana* | 2.625 | Neotropical | M3 | NA | Schwarz (1948) |

**Table S3.** List of evolutionary models fitted to body size (ITD) and centrality metrics (specialization d’, relative degree, betweenness centrality, and closeness centrality). Note that Ornstein-Uhlenbeck is the best fitting evolutionary model for all variables of our data.

| | **Brownian-Motion** | | **Early-Burst** | | **Ornstein-Uhlenbeck** | |
|---|---|---|---|---|---|---|
| **Variables** | **AICc** | **ΔAIC** | **AICc** | **ΔAIC** | **AICc** | **ΔAIC** |
| ITD | 98.79 | 5.33 | 101.12 | 7.66 | 93.46 | 0.00 |
| Specialization d’ | 116.16 | 20.71 | 118.51 | 23.06 | 95.45 | 0.00 |
| Relative degree | 110.95 | 9.92 | 113.41 | 12.38 | 101.03 | 0.00 |
| Betweenness centrality | 132.35 | 24.06 | 134.81 | 26.52 | 108.29 | 0.00 |
| Closeness centrality | -18.13 | 17.46 | -15.79 | 19.80 | -35.59 | 0.00 |

**Table S4.** List of linear models including body size (ITD: intertegular distance) and sampling effort (sampEff: number of studies reporting resin collection for each bee species) as predictors. For each response variable, models are sorted in ascending order by ΔAIC. When more than one model for the same response variable exhibits an ΔAIC < 2, these are considered as equally plausible. Significant p-values for each predictor are pointed with an asterisk.

| Response variable | Model | ΔAIC | AICc | $R^2$ | Predictors | β | Std. Error | p-value |
|---|---|---|---|---|---|---|---|---|
| Specialization d' | ~ITD | 0.00 | 121.89 | 0.1161 | log ITD | 0.7370 | 0.2703 | 0.0089* |
| | ~ITD+sampEff | 0.45 | 122.34 | 0.1312 | log ITD | 0.7464 | 0.2680 | 0.0076* |
| | | | | | sampEff | -0.0912 | 0.0674 | 0.1823 |
| | ~ITD*sampEff | 1.72 | 123.61 | 0.1334 | log ITD | 1.1199 | 0.3237 | 0.0149* |
| | | | | | sampEff | 0.0095 | 0.1165 | 0.9352 |
| | | | | | log ITD:sampEff | -0.1612 | 0.1521 | 0.2947 |
| | ~1 | 4.93 | 126.82 | - | - | -1.9286 | 0.1177 | $2\times10^{-16}$ * |
| | ~sampEff | 5.72 | 127.61 | 0.0088 | sampEff | -0.0863 | 0.0719 | 0.2360 |
| Degree | ~ITD+sampEff | 0.00 | 170.21 | 0.1981 | log ITD | 0.6231 | 0.3813 | 0.1085 |
| | | | | | sampEff | 0.3379 | 0.0984 | 0.0012* |
| | ~sampEff | 0.41 | 170.62 | 0.1718 | sampEff | 0.3432 | 0.0999 | 0.0012* |
| | ~ITD*sampEff | 2.23 | 172.44 | 0.1850 | log ITD | 0.8569 | 0.6521 | 0.1949 |
| | | | | | sampEff | 0.4007 | 0.1729 | 0.0247* |
| | | | | | log ITD:sampEff | -0.0997 | 0.2247 | 0.6590 |
| | ~ITD | 8.87 | 179.08 | 0.0284 | log ITD | 0.6665 | 0.4194 | 0.1180 |
| | ~1 | 9.18 | 179.39 | - | - | -3.3025 | 0.1751 | $2\times10^{-16}$ * |
| Closeness | ~sampEff | 0.00 | -57.52 | 0.2284 | sampEff | 0.0456 | 0.0114 | 0.0002* |
| | ~ITD+sampEff | 2.35 | -55.17 | 0.2124 | log ITD | 0.0036 | 0.0460 | 0.9371 |
| | | | | | sampEff | 0.0455 | 0.0116 | 0.0002* |
| | ~ITD*sampEff | 3.75 | -53.76 | 0.2122 | log ITD | 0.0680 | 0.0795 | 0.3962 |
| | | | | | sampEff | 0.0618 | 0.0201 | 0.0034* |
| | | | | | log ITD:sampEff | -0.0264 | 0.0266 | 0.3251* |
| | ~1 | 11.99 | -45.53 | - | - | -4.1331 | 0.0210 | $2\times10^{-16}$ * |
| | ~ITD | 14.15 | -43.37 | 0.0021 | log ITD | 0.0167 | 0.0522 | 0.7500 |
| Betweenness | ~sampEff | 0.00 | 117.83 | 0.0792 | sampEff | 0.2348 | 0.1185 | 0.0560 |
| | ~ITD+sampEff | 0.74 | 118.57 | 0.0984 | log ITD | -0.6661 | 0.5104 | 0.2012 |
| | | | | | sampEff | 0.2348 | 0.1173 | 0.0539 |
| | ~1 | 1.53 | 119.36 | - | - | -4.2225 | 0.2145 | $2\times10^{-16}$ * |
| | ~ITD*sampEff | 1.79 | 119.62 | 0.1131 | log ITD | 0.3955 | 0.9963 | 0.6941 |
| | | | | | sampEff | 0.4566 | 0.2138 | 0.0407* |
| | | | | | log ITD:sampEff | -0.3467 | 0.2803 | 0.2254 |
| | ~ITD | 2.31 | 120.14 | 0.0162 | log ITD | -0.6661 | 0.5331 | 0.2200 |

**Table S5.** Pearson's correlation matrix between centrality metrics, with p-values in parentheses. Non-significant values ($p > 0.05$) are highlighted in gray. Note that Bluthgen's specialization d' is not significantly correlated with other node-level metrics.

| | Betweenness | Closeness | Degree | Specialization d |
|---|---|---|---|---|
| **Betweenness** | 1 | — | — | — |
| **Closeness** | r = 0.75 (p = 2.44×e$^{-10}$) | 1 | — | — |
| **Degree** | r = 0.40 (p = 3.42×e$^{-3}$) | r = 0.64 (p = 4.40e$^{-7}$) | 1 | — |
| **Specialization d'** | r = -0.09 (p = 0.49) | r = -0.26 (p = 0.06) | r = 0.09 (p = 0.48) | 1 |

### Supplementary Figures

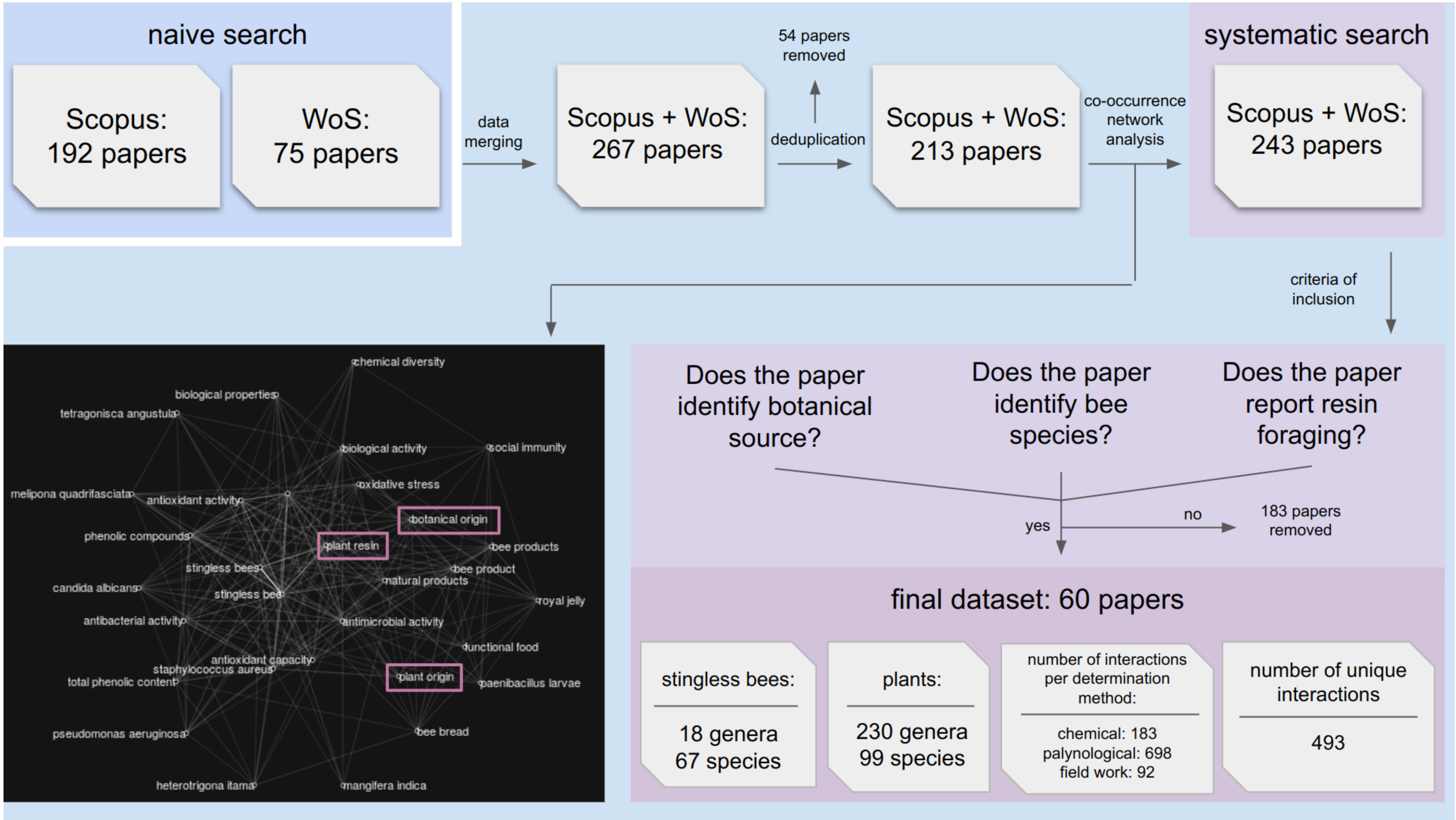


**Figure S1.** Diagram workflow reporting each step of systematic literature search using the co-occurrence network framework by Grames et al. (2019). The keywords “botanical origin”, “plant resin”, and “plant origin” are highlighted in the network, because they were important keywords not originally included in the naive search. Then, they were responsible for enriching the dataset from 213 papers in naive search to 243 in the systematic literature search. After screening each article to check if they pass our criteria of inclusion, our final dataset comprised 60 papers.

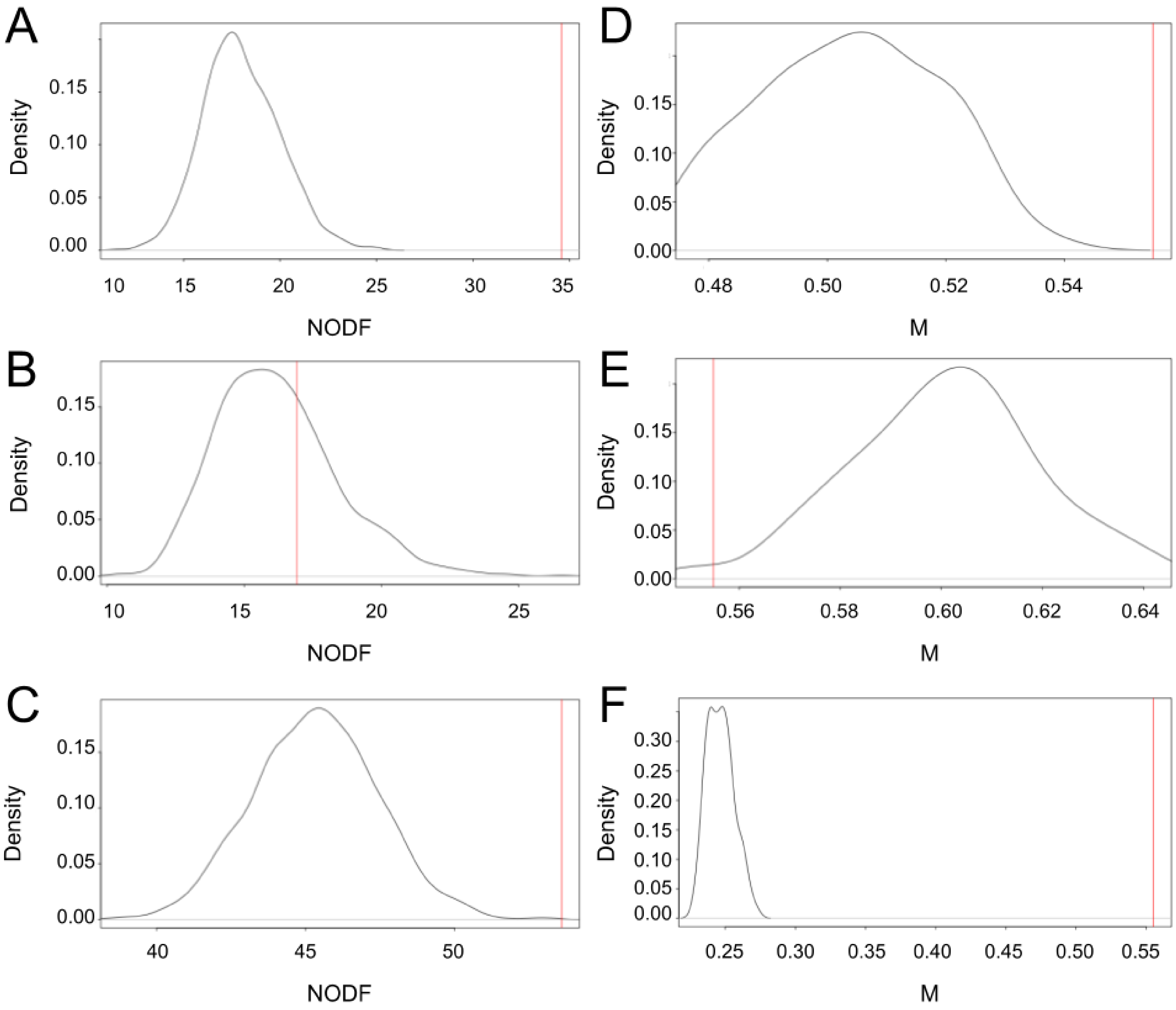


**Figure S2.** Observed NODF values and null models for meta-network assembled from (A) chemical ($NODF_{chemical}$ = 21.99; $NODF_{null\text{-}chemical}$ = 18.01; Z = 8.31; p < 0.001), (B) fieldwork ($NODF_{fieldwork}$ = 16.92; $NODF_{null\text{-}fieldwork}$ = 16.18; Z = 0.33; p = 0.673), and (C) palynological data ($NODF_{palynological}$ = 53.59; $NODF_{null\text{-}palynological}$ = 45.33; Z = 3.83; p < 0.001). Likewise, observed M (modularity) values and null models for meta-network assembled from (D) chemical ($M_{chemical}$ = 0.56; $M_{null\text{-}chemical}$ = 0.50; Z = 3.48; p < 0.001), (E) fieldwork ($M_{fieldwork}$ = 0.55; $M_{null\text{-}fieldwork}$ = 0.61; Z = -2.43; p = 0.98), and (F) palynological data ($M_{palynological}$ = 0.55; $M_{null\text{-}palynological}$ = 0.24; Z = 32.69; p < 0.001).

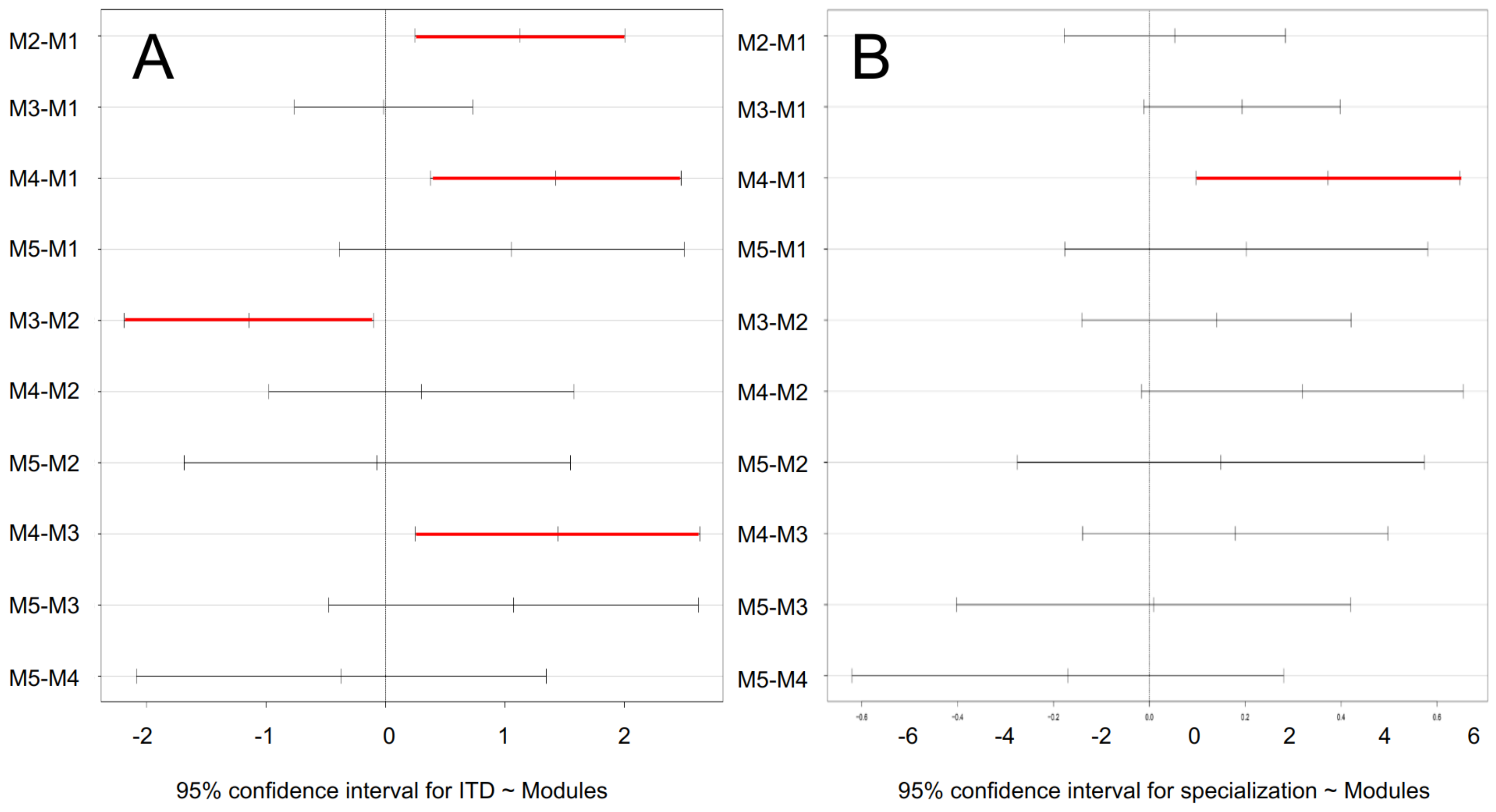


**Figure S3.** Tukey's test with 95% confidence interval testing the difference between (A) body size (ITD: intertegular distance) and modules, and (B) Bluthgen's d specialization and modules. 95% confidence intervals indicating significant difference are colored in red.

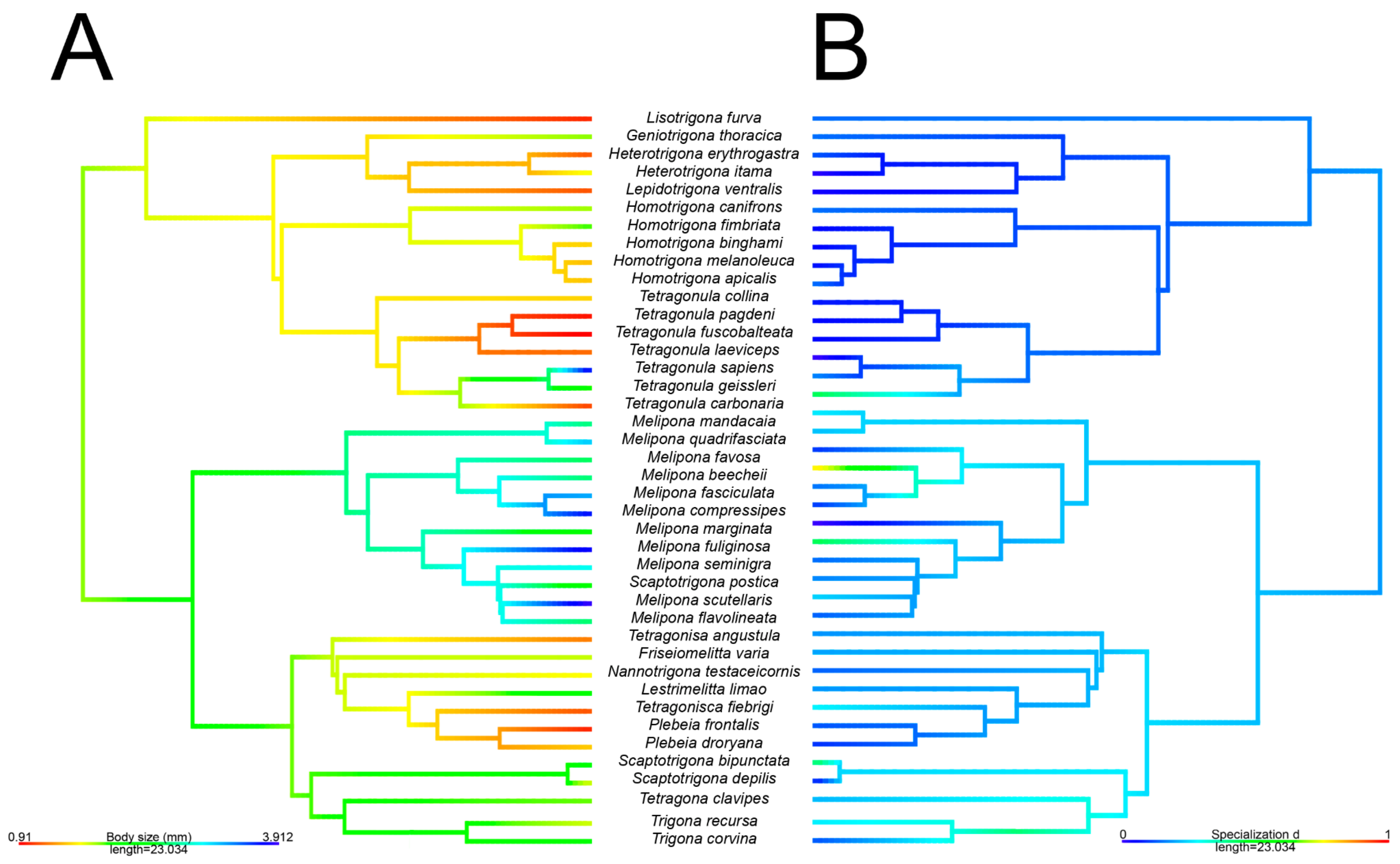


**Figure S4.** Ancestral character state of (A) body size (ITD) and (B) specialization under the Ornstein-Uhlenbeck model using the *contMap* function in 'phytools' (body size: λ = 0.59; p < 0.001, specialization d': λ = 0.19; p = 0.11).

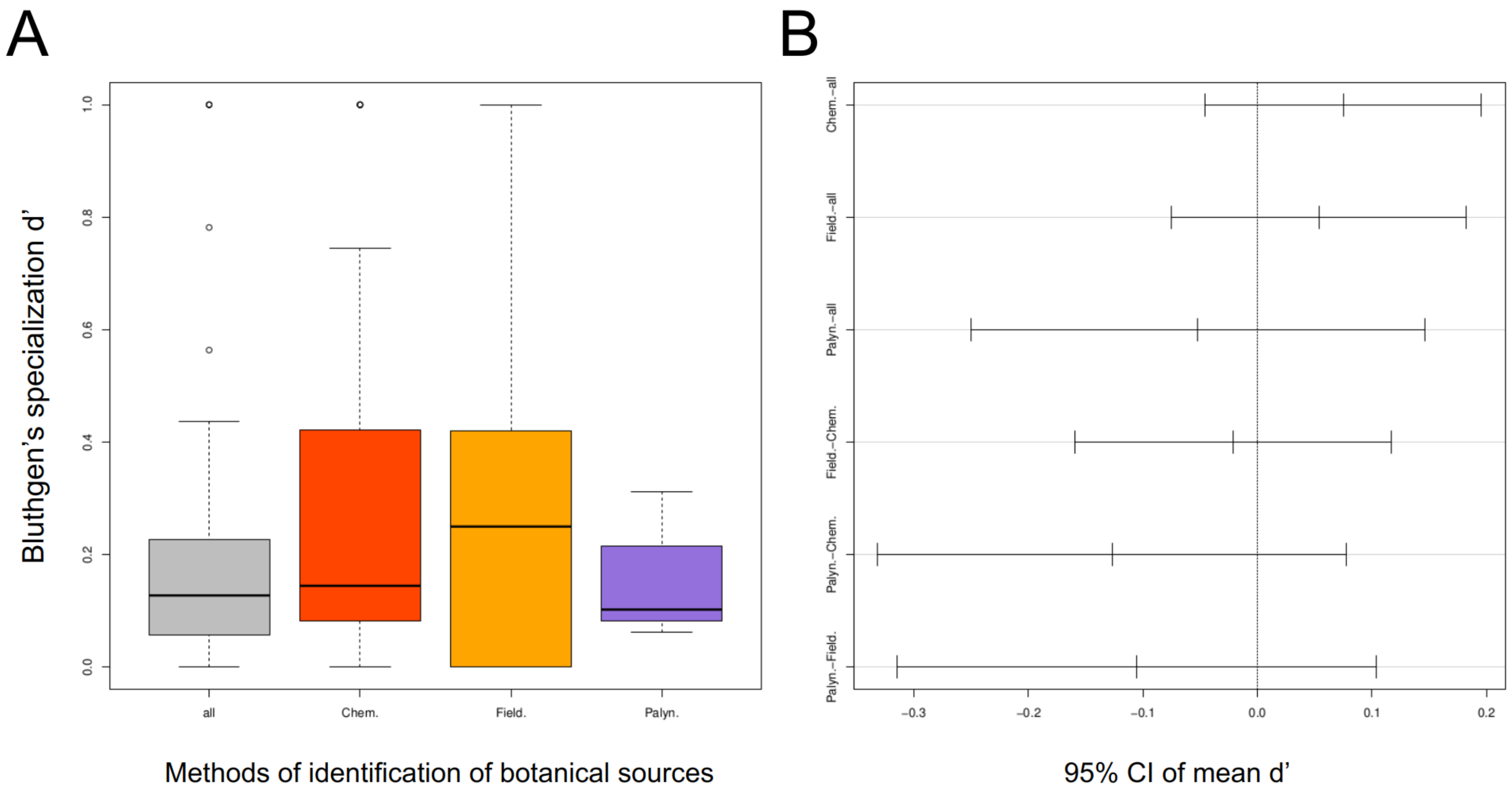


**Figure S5.** Testing whether methods of identification of botanical sources are associated with Bluthgen's specialization d'. (A) Boxplots with ANOVA (F = 1.44; p = 0.23) and (B) Tukey's test with 95% confidence interval suggest no bias from methods to determine botanical sources toward distributions of d'. Abbreviations: chem. = chemical; field. = fieldwork. palyn. = palynological.

### Appendix

#### Glossary of network theory concepts used in this study

**Network-level concepts**

*(I) Disconnected and connected networks*

Ecological networks can be classified in two basic structures: (*a*) disconnected networks comprising a set of isolated components; and (*b*) a fully connected network, commonly with a giant component defined as the set of pathways connecting the highest proportion of nodes in a network (Guimarães 2020). While disconnected networks are characterized by a small average number of interactions per species (low connectivity), connected networks present at least a few species interacting with several other ones (high connectivity; Newman et al. 2001). In the resin foraging network, the first case may happen when a few isolated bee species interact with a subset of plants whose resins are not collected by other bee species (Fig. A1A). On the other hand, the second case happens when all bees and plants have direct or indirect pathways of interactions connecting them (Fig. A1B).

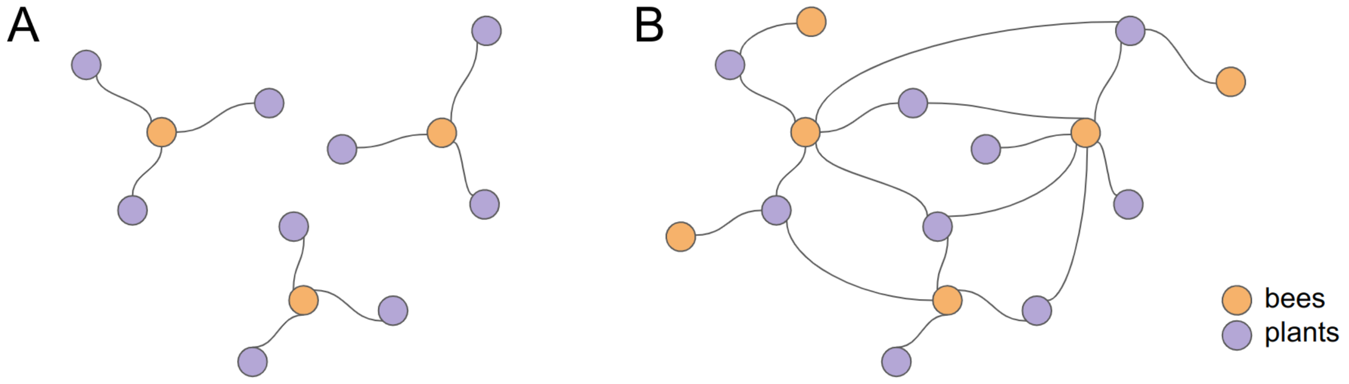


Figure A1: Basic topological structures of bipartite networks. (A) Bipartite disconnected network with a subset of isolated components. (B) Bipartite connected network.

*(II) Modularity*

Modularity is a property that describes the extent to which groups of nodes called modules have a higher probability of being connected to each other than to nodes from other modules (Fortunato and Hric 2016). In ecological networks, modularity emerges when species form more interactions among themselves within a module but fewer interactions between nodes of different modules. Highly connected nodes within a module are called hubs, whereas nodes connecting different modules are called connectors (Fig. A2A; Olessen et al. 2007). Different indexes to detect modularity have been proposed, but the DIRTLPAwd+ algorithm was demonstrated to outperform other methods (Beckett 2016). DIRTLPAwd+ is based on simulations that search for the optimal partition of the network that maximizes the Barber's (2007) modularity index, which is an bipartite version of the Newman's (2006) modularity calculation. It can be mathematically described as:

$$Q=\sum_{m=1}^{N}\left[\frac{L_m}{L}-\left(\frac{K_m^C K_m^R}{L^2}\right)\right]$$

where Q is the modularity index that must be maximized, N is the number of modules, L is the total number of interactions, $L_m$ is the number of links within a module m, $K_m^C$ and $K_m^R$ are the sum of degrees of all bee- and plant-nodes of module m.

*(III) Nestedness*

Nestedness is a property that reflects the magnitude to which interactions of any node form a subset of interactions of all nodes with high degrees (Fig. A2B; Payrató-Borras et al. 2020). We choose the NODF indice (Almeida-Neto et al. 2008) as an operational variable of nestedness, which is defined as:

$$NODF = \frac{\sum_{i<j}^{r} \frac{n_{ij}^{R}}{min(k_i^R, k_j^R)} * \Theta(k_i^R - k_j^R) + \sum_{i<j}^{c} \frac{n_{ij}^{C}}{min(k_i^C, k_j^C)} * \Theta(k_i^C - k_j^C)}{\frac{r(r-1)}{2} + \frac{c(c-1)}{2}}$$

where r is the number of resources (plants); c is the number of consumers (stingless bees); $n_{ij}^R$ and $n_{ij}^C$ are the numbers of shared partners among plants and consumers, respectively; $k_{ij}^R$ and $k_{ij}^C$ are the degrees of resource and consumer, that is, the number of edges connected to each node; Θ is the Heaviside step function that ensures that only pairs of species in which $k_i > k_j$ will contribute to the summation (Felix et al. 2022).

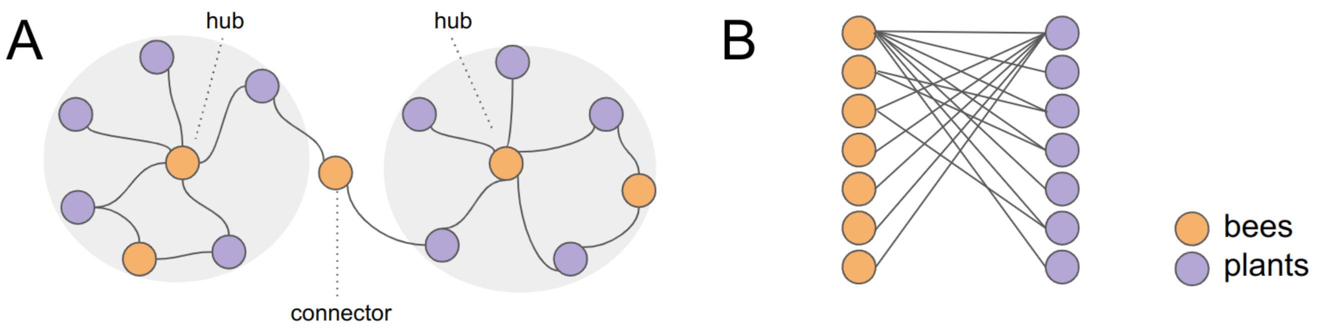


Figure A2: Two common patterns of topological structure in ecological networks. (A) Network depicting two modules, with hubs connecting several nodes within a module and a connector linking the modules. (B) An example of a nested network.

**Node-level metrics**

The node-level metrics employed in this study are centrality metrics, that is, metrics measuring the relative importance of a node to the topology of its network. Although some centrality metrics might be correlated (see Supplementary Table S4), we employed four centrality metrics because each one is focused on different aspects of a node importance and thus may be interpreted differently in an ecology scope.

*(I) Betweenness centrality*

Betweenness centrality is a node-level metric associated with the magnitude to which a node is a bridge between different regions of a network (Fig. A3A). It is calculated as the division between the number of shortest paths (i.e., geodesic distance, the minimum number of edges between two nodes) in which a given species is present in the empirical network and the number of potential geodesic distances in a simulated star network (Freeman 1977). In ecological networks, betweenness centrality may be interpreted as the magnitude of a species in binding different guilds within the network (Mello et al. 2019).

*(II) Bluthgen's specialization d'*

Bluthgen's d specialization is a node-level metric associated with the selectiveness of the set of interactions performed by a bee in relation to the interactions made by all other bee species in the network. Blüthgen et al. (2007) originally named it as Kullback-Leibler distance (d') based on relative entropy and information theory (Kullback and Leibler 1951). To calculate d', interaction data must be written as an incidence matrix comprising rows i (plants) and columns j (stingless bees), within which each cell is given as $a_{ij}$ (zero or one for binary networks or the frequency of interactions for weighted networks). The sum of interactions for each bee species is $A_j$, whereas the sum of interactions for each plant is $A_i$. The total number of interactions in the network is m. Bluthgen's specialization d' for a species i ($d_i$) is calculated as:

$$d_i = \sum_{j=1}^{c} \left( p'_{ij} \cdot \ln \frac{p'_{ij}}{q_j} \right)$$

where $p'_{ij}$ is the proportion of the number of interactions ($a_{ij}$) in relation to the respective row ($A_i$), $q_j$ is the proportion of all interactions by bee j ($A_j$) in relation to the total number of interactions (m). Therefore, d' compares the distribution of interactions with each partner ($p'_j$) to its overall availability in the system ($q_j$). In our case, a high specialization occurs when a bee species collects resins from a set of plant species that are not strongly collected by other bee species (Fig. A3B).

*(III) Closeness centrality*

Closeness centrality quantifies the magnitude to which a node can access all other nodes in connected networks. Hence, the more central the node, the closer it is to all nodes (Farahani et al. 2019). Nodes with a high closeness value have shortest paths (geodesic distances) to all other nodes (Fig. A3C). In ecology, this metric may be interpreted as a proxy for niche overlap (Mello et al. 2015).

*(IV) Relative degree*

Relative degree is the number of node's neighbors (Costa 2004) scaled by the total number of nodes in the network (Fig. A3D). In ecology, it might be interpreted as the fundamental niche breadth (Mello et al. 2015).

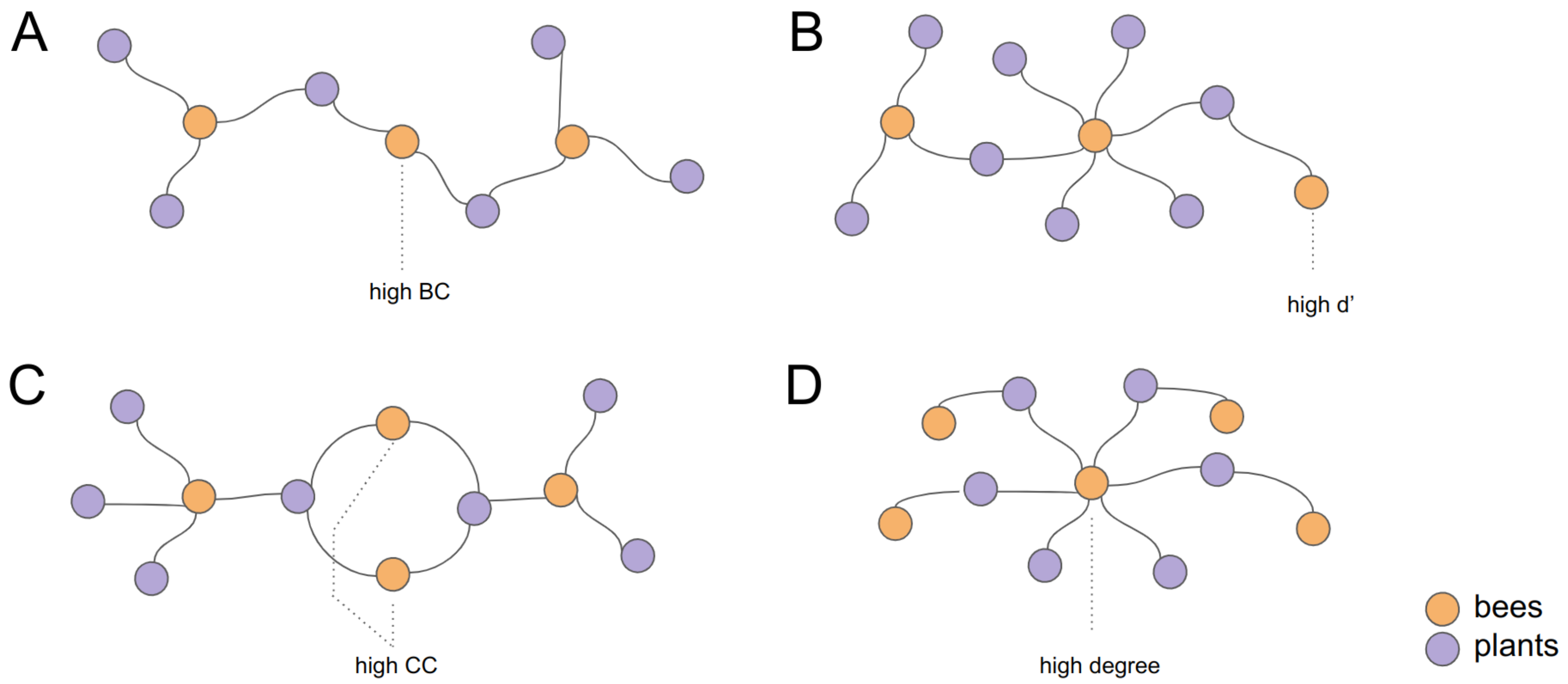


Figure A3: Centrality-metrics employed in this study. (A) Betweenness centrality (BC). (B) Bluthgen's specialization d'. (C) Closeness centrality (CC). (D) Relative degree.
